# Differential Effects of Mutations of Popeye Domain Containing Proteins on Heteromeric Interaction and Membrane Trafficking

**DOI:** 10.1101/2022.10.13.511879

**Authors:** Alexander H. Swan, Roland F.R. Schindler, Marco Savarese, Isabelle Mayer, Susanne Rinné, Felix Bleser, Anne Schänzer, Andreas Hahn, Mario Sabatelli, Francesco Perna, Kathryn Chapman, Mark Pfuhl, Alan C. Spivey, Niels Decher, Bjarne Udd, Giorgio Tasca, Thomas Brand

**Affiliations:** National Heart and Lung Institute, Imperial College London, London, United Kingdom; Department of Chemistry, Imperial College London, London, United Kingdom; Assay Biology, Domainex Ltd, Cambridge, CB10 1XL, United Kingdom; Department of Medical Genetics, Medicum, University of Helsinki, Helsinki, Finland; Institute for Physiology and Pathophysiology, Vegetative Physiology, Philipps-University of Marburg, Marburg, Germany; Institute of Neuropathology, Justus Liebig University Giessen, 35392 Giessen, Germany; Department of Child Neurology, Justus Liebig University Giessen, 35392 Giessen, Germany; Department of Neurology, Universitá Cattolica del Sacro Cuore, Rome, Italy; Dipartimento di Scienze Cardiovascolari, Fondazione Policlinico Universitario A. Gemelli IRCCS, Rome, Italy; School of Cardiovascular Medicine and Sciences and Randall Centre, King’s College London, Guy’s Campus, London SE1 1UL, United Kingdom; Folkhälsan Research Center, University of Helsinki, Helsinki, Finland; Unità Operativa Complessa di Neurologia, Fondazione Policlinico Universitario A. Gemelli IRCCS, Rome, Italy

**Keywords:** Popeye domain containing protein, Popeye domain, limb-girdle muscular dystrophy, membrane targeting, protein interaction

## Abstract

**Background:** The Popeye domain containing (POPDC) genes encode sarcolemma-localised cAMP effector proteins. Mutations in *BVES (POPDC1)* and *POPDC2* have been associated with limb-girdle muscular dystrophy and cardiac arrhythmia. Muscle biopsies of affected patients display impaired membrane trafficking of both POPDC isoforms.

**Methods:** Biopsy material of patients carrying mutations in *BVES* were immunostained with POPDC antibodies. The interaction of POPDC proteins was investigated by co-precipitation, proximity ligation, bioluminescence resonance energy transfer and bimolecular fluorescence complementation. Site-directed mutagenesis was utilised to map the domains involved in protein interaction.

**Findings:** Patients carrying a novel homozygous variant, *BVES* (c.547G>T, p.V183F) displayed only a skeletal muscle pathology and a mild impairment of membrane trafficking of both POPDC isoforms. This is in contrast to variants such as *BVES* p.Q153X or *POPDC2* p.W188X, which were associated with a greater impairment of membrane trafficking. Co-transfection analysis in HEK293 cells revealed that POPDC proteins interact with each other through a helix-helix interface located at the C-terminus of the Popeye domain. Site-directed mutagenesis of an array of ultra-conserved hydrophobic residues demonstrated that some of them are required for membrane trafficking of the POPDC1-POPDC2 complex.

**Interpretation:** Mutations in POPDC proteins that cause an impairment in membrane localisation affect POPDC complex formation while mutations which leave the protein interaction intact likely affect some other essential function of POPDC proteins.

**Funding:** This study was funded by an EPSRC/British Heart Foundation co-funded Imperial Institute of Chemical Biology (ICB) Centre for Doctoral Training (CDT) PhD studentship (EP/S023518/1), a project grant of the British Heart Foundation (PG19/13/34247) and the Deutsche Forschungsgemeinschaft (DE1482/9-1).

**Research in Context:** *Evidence before this study:* Several biallelic missense and nonsense variants in *BVES (POPDC1)* have been described and are associated with heart and skeletal muscle disease. Skeletal muscle biopsies of homozygous carriers of these variants display a loss of sarcolemmal localisation of POPDC1 and POPDC2.

*Added value of this study:* We demonstrate that POPDC1 and POPDC2 form a heteromeric complex and that complex formation is required for plasma membrane trafficking of POPDC proteins. Transfection of different disease variants in HEK293 cells replicates their defective membrane targeting observed in biopsy material. Structural modelling and site-directed mutagenesis identifies an interface of strongly conserved hydrophobic residues in POPDC proteins, which likely mediate the interaction of POPDC proteins.

*Implications of all the available evidence:* These data provide novel insight into the membrane targeting requirements of POPDC proteins. We recommend testing the membrane targeting properties of any novel variant in POPDC isoforms using a newly developed co-transfection assay in HEK293 cells to characterise its pathogenicity. Our novel insight into the requirement of heterodimerization for proper membrane targeting may also offer novel opportunities to treat patients carrying mutations in POPDC proteins.

## Introduction

The Popeye domain containing (POPDC) gene family consists of three family members, blood vessel epicardial substance (*BVES*, also known as *POPDC1*), *POPDC2*, and *POPDC3*.^1,2^ POPDC genes encode transmembrane proteins, which are abundantly expressed in the sarcolemma of cardiac and skeletal muscle cells.^3^ POPDC proteins consist of a short extracellular amino-terminus, which is subject to N-glycosylation followed by three transmembrane domains.^4^ The cytoplasmic part of the protein consists of the Popeye domain and a carboxy-terminus, which is isoform-specific and of variable length.^3^ The Popeye domain binds 3’,5’-cyclic adenosine monophosphate (cAMP) with high affinity and specificity.^5^ In the heart, POPDC1 and POPDC2 are expressed in cardiac myocytes and both isoforms display high expression levels in the cardiac conduction system (CCS).^1,5–7^ Consistent with their CCS expression, a near identical stress-induced sinus node bradycardia was observed in *Bves* and *Popdc2* knockout (KO) mice.^5^ Enhanced vulnerability of the mutant heart in response to ischemia-reperfusion and impaired skeletal muscle regeneration after injury have also been described for the *Bves* KO mutant.^8,9^ Similarly, cardiac arrhythmia and muscular dystrophy are present in the zebrafish *bves* KO mutant and *popdc2* morphants.^10,11^ POPDC proteins function as a novel class of cAMP effector proteins ^3^ and interactions with other proteins involved in cAMP signalling such as phosphodiesterase 4 (PDE4) and adenylyl cyclase 9 (AC9) have recently been described.^12,13^

A sizable number of patients who carry pathogenic variants in POPDC genes have been identified and suffer from heart and/or muscle disease.^11,14–17^ Patients carrying *BVES* mutations develop a recessive form of limb-girdle muscular dystrophy (LGMDR25), with most patients also suffering from cardiac arrhythmia (sinus bradycardia, sinus tachycardia, or atrioventricular (AV)-block).^11,17–21^ Some patients display structural changes in the heart including thickening of the septum and dilated cardiomyopathy.^19,21^ Heart disease with no apparent skeletal muscle involvement has been described in one family.^20^ It can be concluded that the pathology caused by *BVES* mutations displays high variability regarding the age of onset, phenotype severity and affected organs. Several reports described a loss of sarcolemmal expression of both POPDC1 and POPDC2 when skeletal muscle biopsies of patients carrying mutations in *BVES* were investigated.^11,17,19^

In this study, we report a novel recessive mutation in *BVES* (c.547G>T, p.V183F) which has been discovered in two unrelated patients suffering from limb-girdle muscular dystrophy with no cardiac involvement. In contrast to the loss of sarcolemmal expression described for other *BVES* variant cases, expression of POPDC1 and POPDC2 was only weakly diminished in biopsy material of both patients. Currently, it is unclear what determines membrane trafficking of POPDC proteins. We established by co-transfection analysis in HEK293 cells that co-expression of both POPDC1 and POPDC2 was required for proper membrane localisation. POPDC proteins undergo heteromeric complex formation as demonstrated by proximity ligation and bioluminescence resonance energy transfer (BRET) analysis. Membrane trafficking is controlled by the formation of an interface between α-helices located at the carboxy-terminus of the Popeye domain of POPDC1 and POPDC2. Modelling identified an array of ultra-conserved hydrophobic residues in both isoforms. Support for their possible involvement in mediating membrane trafficking of POPDC1 and POPDC2 was obtained by site-directed mutagenesis. We propose a model to explain the differential effect of different *BVES* and *POPDC2* variants on membrane trafficking.

## Methods

### Study subjects, clinical and molecular examinations

Two unrelated patients affected by a primary muscle disorder were genetically investigated. Genomic DNA was isolated from their blood cells using standard techniques. DNA samples of patients one (PT1) and two (PT2) were sequenced using a targeted gene panel (Myocap) ^22^ and a clinical exome filtered for 206 myopathy-associated genes respectively. Raw NGS data were analysed using a standard pipeline. *BVES* variants, described on transcript NM_001199563, were confirmed by PCR and Sanger sequencing (primers available on request) and their segregation analysis was performed on the available family members. Patients and their unaffected relatives provided written informed consent. The study was approved by the Ethics Review Board of Helsinki University Hospital (number 195/13/03/00/11) according to the Declaration of Helsinki. The work on the patient carrying the Q153X variant was approved by the ethical committee of the University of Giessen (AZ07/09, AZ258/16). Written informed consent was obtained from the parents of the patient. The study was conducted in accordance with the principles of the Declaration of Helsinki.

### Animal work

All experimental mouse work was ethically approved by the Animal Welfare and Ethical Review Board of Imperial College London and licensed by the United Kingdom Home Office (PPL P2960EB2F) and conformed to the guidelines from the Directive 2010/63/EU of the European Parliament on the protection of animals used for scientific purposes. All experiments were performed using age-matched mice. C57BL/6J mice (RRID: IMSR_JAX:000664) were purchased from Harlan UK. The mice were housed in standard cages with a 12-h light/12-h dark cycle at 22 to 24 °C and *ad libitum* access to food and water. A *Popdc2* p.W188X knockin (KI) mutation was generated by homologous recombination in embryonic stem cells.^14^ *Popdc1*/*Popdc2* double KO animals were generated by crossing *Popdc1* and *Popdc2* KO animals.^5^ Both lines were at least ten times backcrossed with C57Bl6J mice and were kept subsequently in a homozygous state. Heart and skeletal muscle tissue of mutant and wild-type mice was obtained after euthanasia. The animal study using *Xenopus* toads was approved by the Ethics Committee of the Regierungspräsidium Giessen (protocol code V54-19c 20 15 h 02 MR 20/28 Nr. A23/2017, approved on the 12.02.2018).

### Immunostaining

Muscle biopsies from patients and mouse skeletal muscle were processed according to standard procedures.^11,23^ Sections were mounted on Superfrost glass slides (Thermo Fisher Scientific) and subjected to immunohistochemistry using the following primary antibodies: POPDC1 (HPA018176, Sigma-Aldrich), POPDC2 (HPA024255, Sigma-Aldrich), α-sarcoglycan (SGCA, NCL–α-SARC, Leica Biosystems). For the detection of primary antibodies, the following secondary antibodies were employed: Alexa Fluor 488–conjugated donkey anti-rabbit (A21206, Invitrogen) and Alexa Fluor 555–conjugated donkey anti-mouse (A31570, Invitrogen). For counterstaining, DAPI (Calbiochem) was employed. Sections were imaged using a Zeiss LSM 780 AxioObserver inverted confocal laser scanning microscope, with a plan-apochromat 20X/0·8 M27 objective (Zeiss). Three channels were used during image acquisition: 405_ex_/410-495_em_ nm for DAPI, 488_ex_/489-552_em_ nm for Alexa Fluor 488, 543_ex_/548-697_em_ nm for Alexa Fluor 544. 3-15 images were taken for each analysis group.

### Image analysis of skeletal muscle fibres

Images of the immunohistochemically stained skeletal muscle sections were processed using FIJI.^24^ The SGCA channel was used to automatically produce outlines of individual muscle fibre cross-sections through the use of a threshold limit. Dilation of the fibre outlines was used to enclose the sarcolemma, with the cytoplasm defined as the inner area of each fibre. The average intensity of Alexa Fluor 488 (POPDC1 or POPDC2) and Alexa Fluor 544 (SGCA) fluorescence within the sarcolemma and cytoplasm of each fibre was then determined. The SGCA-normalised sarcolemmal expression of POPDC1 and POPDC2 was determined by dividing the Alexa Fluor 488 signal within the sarcolemma compartment by that of Alexa Fluor 544. Additionally, the cross-sectional area of each fibre was recorded.

### Cell Culture

HEK293 (DSMZ, RRID: CVCL_0045) and COS-7 (DSMZ, RRID: CVCL_0224) were cultured in Dulbecco’s modified Eagle’s medium (Sigma Aldrich) supplemented with 10% (v/v) foetal bovine serum (Merck). HEK293 and COS-7 cells were transiently transfected using Lipofectamine 2000 (Invitrogen) or calcium phosphate (Promega). Cells were incubated for 24-48-hrs post-transfection before use.

### Cloning Procedures

Full length human *POPDC1* and *POPDC2* cDNAs were inserted into the pECFP-N1/pEYFP-N1 plasmid, to append a C-terminus ECFP/EYFP tag. The clinically identified POPDC variants, and the aspartic acid scanning mutations of hydrophobic residues in the αC-helix, were introduced into these constructs using the Q5 site-directed mutagenesis kit (NEB). The oligonucleotide primer sequences used in the site-directed mutagenesis reactions are listed in Supplemental Table 1. For NanoBRET analysis, human *POPDC1* and *POPDC2* cDNA sequences were cloned into the pFC14K or pFC32K plasmids (Promega), which contain C-terminus sequences for HaloTag and NanoLuc tags, respectively, using the SgfI and EcoICRI restriction sites. For bimolecular fluorescence complementation (BiFC), split Venus VN155 and VC155 tags (kindly provided by Carmen Dessauer, University of Texas) were ligated into pECFP-N1 or pEYFP-N1 plasmids containing full-length wild-type and mutant POPDC cDNA sequences using NotI and BamHI restriction sites.

### Quantitative BRET

Type-1 quantitative bioluminescence resonance energy transfer (qBRET) experiments utilised the NanoBRET platform (Promega) and followed the general type-1 qBRET protocol reported by Felce et al.^25^ The assay was performed in HEK293 cells transiently expressing POPDC1 and POPDC2 constructs which possessed a C-terminal NanoLuc luciferase or HaloTag domains. 100 nM of HaloTag-618 dye (Promega) was added to the cells 24-hrs before BRET measurement, with an equal number of cells receiving only DMSO to enable the determination of background BRET, which was subtracted from final BRET values. Cells were placed in white 96-well tissue culture plates, immersed in OptiMEM I reduced serum media supplemented with 4% (v/v) foetal bovine serum, 24-hrs before BRET measurement. BRET was measured using a Lumistar Optima luminometer (BMG Labtech) 5-mins after the addition of furimazine NanoLuc substrate (Promega). A range of expression ratios of the NanoLuc and HaloTag containing constructs within the cells was achieved by varying the proportion of each plasmid during transfection, while keeping the total amount constant. Actual expression levels of the NanoLuc- and HaloTag-fused constructs were determined by measuring the total luminescence and HaloTag-618 fluorescence from the cells, respectively. The total expression levels of POPDC isoforms within the cells was determined by summing the normalised NanoLuc luminescence and HaloTag-618 fluorescence. The BRET curves produced were compared to ideal curves for monomers, dimers and other stoichiometries to determine the likely POPDC complex stoichiometry according to the method reported by Felce et al.^25^

### Co-expression of POPDC isoforms in HEK293 cells

HEK293 cells transiently expressing POPDC1 and POPDC2 constructs tagged at the C-terminal with either ECFP or EYFP were incubated with 0.5% (v/v) CellBrite Red solution (Biotium) containing 1,1’-dioctadecyl-3,3,3’,3’-tetramethylindo-dicarbocyanine (DiD) for 12-mins at 37 °C to stain the plasma membrane. Cells were washed with PBS before fixation with PFA and stained with Hoechst-33342. Cells were imaged using a LSM 780 AxioObserver inverted confocal laser scanning microscope (Zeiss), with a plan-apochromat 63X/1.40 oil objective (Zeiss). Four channels: 405_ex_/410-452_em_ nm, 458_ex_/463-516_em_ nm, 514_ex_/519-621_em_ nm and 633_ex_/636-735_em_ nm, were used to image the Hoechst-33342, POPDC1-ECFP, POPDC2-EYFP and DiD, respectively. Cells were imaged in poly-L-lysine coated 8-well microscope slides with a D263 M Schott glass, No. 1.5H, 170 μm ± 5 μm glass coverslip base (Ibidi) immersed in PBS.

Images were analysed using FIJI. The plasma membrane of the HEK293 cells was manually outlined using the DiD channel as a guide, followed by dilation to encompass the entire plasma membrane. The area within the inner edge of the plasma membrane boundary up to the nucleus, as highlighted by Hoechst-33342, was defined as the cytoplasm. Background fluorescence was subtracted from all images before analysis. The intensity of ECFP and EYFP fluorescence intensity with each compartment was then determined to find the concentration of POPDC1 and POPDC2, respectively.

### Proximity Ligation Assay

A Duolink® proximity ligation assay (PLA, Sigma-Aldrich) was employed using anti-POPDC1 (Santa Cruz) anti-POPDC2 (Atlas) antibodies on heart sections of wild-type and *Bves/Popdc2* double KO mutants according to the standard protocol. Staining with wheat germ agglutinin and DAPI was used to visualize the sarcolemmal and nuclear compartments, respectively.

### Western Blot and Co-Immunoprecipitation Analysis

Cells expressing POPDC1-CFP and/or POPDC2-FLAG, or POPDC1-CFP and/or POPDC3-FLAG were lysed 24-hrs post-transfection using 4 M urea and 10% (w/v) SDS without a reducing agent. Cell lysates were sonicated and centrifuged for 30-mins at >16,000 g. The cleared lysate was incubated at 37 °C for 30-mins and subjected to Western blot analysis using an anti-GFP antibody (Abcam) to detect POPDC1-CFP containing complexes. Ventricles of wild-type and *Popdc2* KO mutant mice were excised, snap-frozen in liquid nitrogen and pulverised with a pre-cooled pestle and mortar. The tissue was lysed using a 1% (v/v) Triton X-100 based lysis buffer followed by sonification. The lysates were centrifuged for 30-mins at >16,000 g. Equal protein concentrations were used across all samples. Anti-Popdc2 antibodies (Sigma Aldrich) were incubated with the cleared lysate overnight. Antibodies were captured using Protein A agarose, which were centrifuged, washed then resuspended in NuPAGE LDS Sample Buffer (Invitrogen) and incubated at 96 °C for 5-mins. After removal of the remaining agarose by centrifugation the sample was supplemented with NuPAGE Sample Reducing Agent (Invitrogen) and analysed by Western blotting. COS-7 cells transiently expressing POPDC constructs with the appropriate epitope tag were lysed using a 1% (v/v) Triton X-100 based lysis buffer supplemented with cOmplete protease inhibitor cocktail (Roche). Lysates were subjected to Co-IP using the ProFoundTM c-Myc Tag IP/co-IP Kit (Thermo Scientific) or the Pierce HA Tag IP/Co-IP Kit (Thermo Scientific), following the manufacturer’s protocols. The antibody conjugated agarose beads were washed, resuspended in sample buffer and analysed by Western blotting.

### Measurement of TREK-1 current

*Xenopus laevis* were maintained and oocytes isolated under standard conditions according to established protocols. Capped cRNA transcripts were synthesized *in vitro* using the mMessage mMachine T7 transcription kit (Ambion). The cRNAs were purified and photometrically quantified. cRNA coding for human TREK-1c alone or together with mouse Popdc1 or mouse Popdc2 were injected into *Xenopus laevis* oocytes. Oocytes were incubated at 19 °C for 48-hrs in ND96 solution containing 96 mM NaCl, 2 mM KCl, 1 mM MgCl_2_, 1·8 mM CaCl_2_, and 5 mM HEPES (pH 7·5) supplemented with 50 mg/l gentamicin and 275 mg/l sodium pyruvate. For experiments with elevated cAMP levels, 25 mM theophylline was supplemented to the storage solution, directly following the cRNA injection. Two-microelectrode voltage-clamp measurements were performed with a Turbo Tec-10 C amplifier (npi, Tamm). The oocytes were placed in a small-volume perfusion chamber and superfused with ND96 solution. Micropipettes were made from borosilicate glass capillaries GB 150TF-8P (Science Products) and pulled with a DMZ-Universal Puller (Zeitz). The resistance of the recording pipettes was 0·5-1·5 MΩ when pipettes were filled with 3 M KCl solution. TREK-current was measured using a voltage step protocol from a holding potential of −80 mV. A first test pulse to 0 mV of 1 s duration was followed by a repolarizing step to −80 mV for 1 s, directly followed by another 1 s test pulse to +40 mV. The sweep time interval was 10 s. Current amplitudes were analysed at +40 mV. Since current amplitudes varied from one batch of oocytes to the next, currents were normalized to TREK-1c WT current amplitudes of the respective batch and recording day.

### Bimolecular fluorescence complementation (BiFC)

POPDC1 and POPDC2 were tagged at the C-terminal with split Venus domains VC155 or VN155, respectively, and expressed in HEK293 cells. Co-transfection of POPDC1-VN155 and POPDC2-VN155 was used as a negative control. All cells were also transfected with pmRFP-N1 as an internal control for transfection efficiency. Cells were fixed using PFA and stained with Hoechst-33342. The cells were imaged using an Axio Observer inverted confocal laser scanning microscope (Zeiss) using a 10X objective (Zeiss). Three channels were used during image acquisition: 405_ex_/410-503_em_ nm for Hoechst-33342, 514_ex_/516-587_em_ nm for reformed Venus and 543_ex_/582-754_em_ nm for mRFP. Images were analysed using FIJI. All cells expressing above-background levels of mRFP fluorescence were selected for analysis from each image, thus excluding non-transfected cells. The median Venus fluorescence from these cells was determined, disregarding the nuclei as defined by Hoechst-33342 staining. The Venus signal from each image was normalised to mRFP fluorescence and the average from each set of images found.

### Sequence alignments and structural models of POPDC protein

For the sequence alignment shown in Figure 1b, different vertebrate POPDC1 and invertebrate POPDC homologues were identified by BLAST. The sequences used for the alignment have the following accession numbers at the NCBI protein database (https://www.ncbi.nlm.nih.gov/protein): *Homo sapiens* (AAH40502.2), *Mus musculus* (NP_077247.1), *Monodelphis domestica* (XP_016286698), *Ornithoryhnchus anatinus* (XP_028903352.1), *Gallus gallus* (NP_001001299), *Xenopus laevis* (AF527799_1), *Danio rerio* (NP_001244093.1), *Strongylocentrotus purpuratus* (XP_003723894.2), *Ciona intestinalis* (XP_002127439.1), *Aplysia californica*: XP_012939248.2, *Capitella teleta* (ELT88986). AlphaFold Protein Structure Database ^26,27^ models of human POPDC1 and POPDC2 were used for modelling purposes and were analysed using Chimera X.^28^ Possible steric clashes caused by the V183F mutation were predicted using the Dynameomics rotamer library within ChimeraX.^29^ Overlays of CAP and Popeye domain protein structures were created using the Matchmaker tool in ChimeraX. Predictions of cAMP binding to the Popeye domain were made using the Phyre2 and 3DLigandSite servers.^30,31^

**Figure 1.**
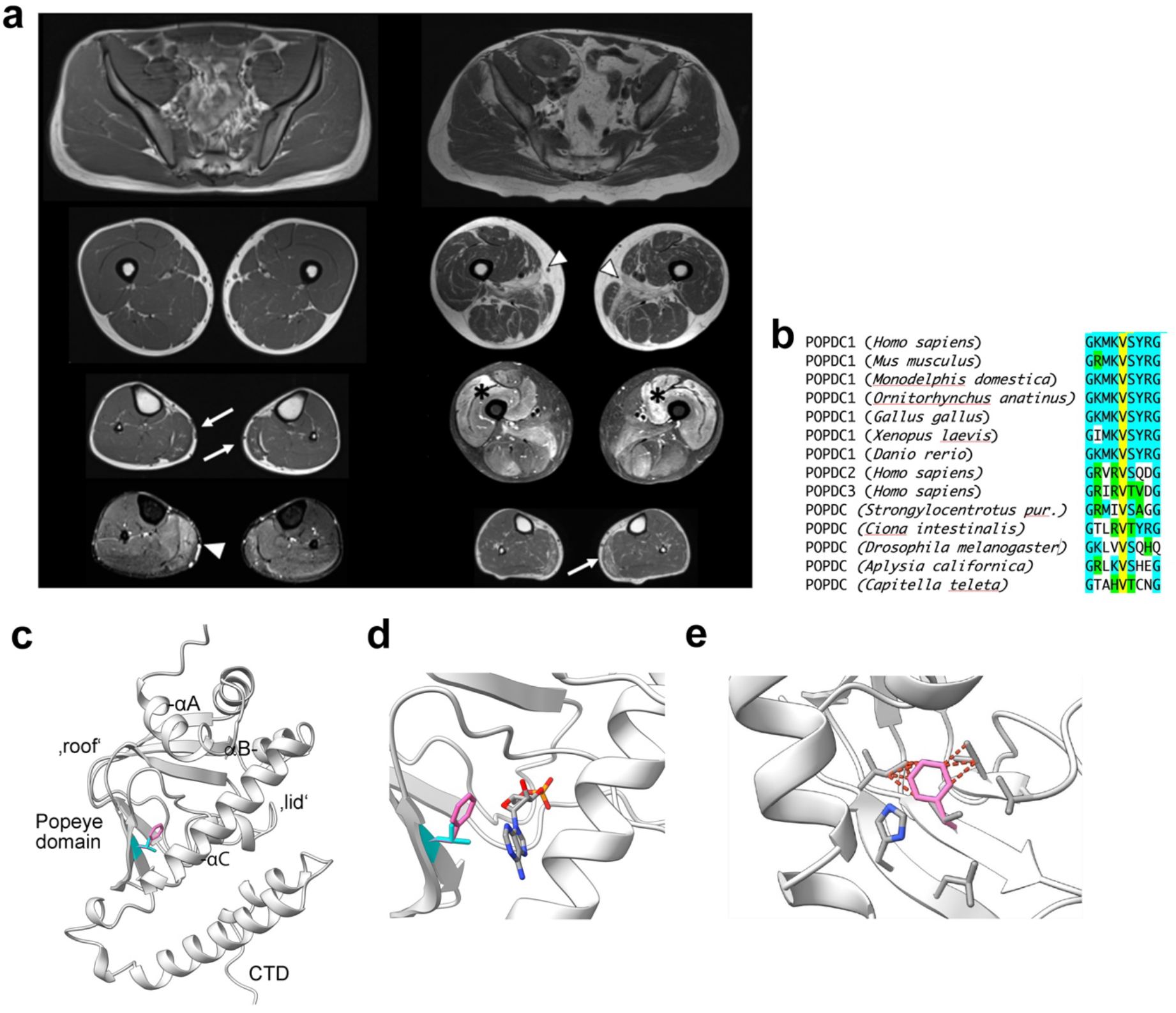
Two non-related patients carrying a *BVES p*.*V183F* variant and suffering from LGMDR25. (**a**) Axial muscle MRI images of patient 1 (PT1, left column) and patient 2 (PT2, right column). PT1 scanned at age 17 displaying fatty replacement and hypotrophy of the *gastrocnemius medialis* (arrows), with hyperintensity on T2-STIR images (arrowheads). PT2 scanned at age 50 displaying, in addition to changes in the *gastrocnemius medialis* (arrows), also advanced fatty replacement of *adductor longus* more than *adductor magnus* (arrowheads) and diffuse T2-STIR hyperintense lesions in the thigh, more evident in the anterior compartment (asterisks). (**b**) Sequence alignment of part of the Popeye domains of vertebrate POPDC1, POPDC2 and POPDC3 and invertebrate POPDC proteins. Colour code: V183 (yellow), conserved (turquois) and similar (green) residues. (**c**) Model of the Popeye domain of POPDC1 with the position of V183 (cyan) and the mutant V183F (pink) residues highlighted. The position of the phenylalanine side chain was determined using the Dynameomics rotamer library.^29^ (**d**) The position of the V183F mutation relative to the predicted cAMP binding site as determined using the 3DLigandSite server.^31^ (**e**) A model showing the possible steric clashes between the side chain of V183F (pink) with other residues of the β-folds of the Popeye domain as predicted by the Dynameomics rotamer library.^29^

### Statistics

The absolute, or SGCA-normalised, expression levels of POPDC1 and POPDC2 within the sarcolemma and cytoplasm of individual muscle fibres were normalised to the median values from matched control fibres, which were set to equal 1. The normalised median and associated 95% CI limits of POPDC1 and POPDC2 expression levels in the patient or KI mutant fibres were determined, then compared using a Mann-Whitney test.

The median difference and 95% CI limit in the ratio of POPDC1-ECFP and POPDC2-EYFP constructs at the plasma membrane versus cytoplasm in HEK293 cells was determined and normalised to the median value for cells expressing both wild-type POPDC1 and POPDC2 constructs. Absolute changes in POPDC1 and POPDC2 expression in each compartment were also recorded. The expression levels between single and double wild-type expression groups were analysed using a Mann-Whitney test, while the effect of the various POPDC mutations, compared to wild-type, were assessed using a Kruskal-Wallis test followed by Dunn’s test.

During the BiFC assay the BiFC signal was defined as the median Venus signal normalised to median mRFP emanating from all transfected cells within each image. The average BiFC signal between the different expression groups were further normalised to the wild-type POPDC pair, set to 1, and compared using a One Way ANOVA followed by Dunnett’s test.

## Results

### A novel POPDC1 p.V183F variant leads to mild loss of POPDC1 and POPDC2 at the sarcolemma

We report here a novel variant in *BVES* (c.547G>T, p.V183F), which has been identified in two unrelated patients. The variant was present in homozygosity in both patients, while the unaffected family members were heterozygous or wild type. The variant is not expected to cause a mis-splicing (SpliceAI max score 0·02) and is classified in VarSome (https://varsome.com/) as a variant of unknown significance (VUS). However, its identification in two unrelated patients with a similar phenotype suggests that there is sufficient evidence for the variant to be classified as VUS/likely pathogenic.

Patient 1 (PT1) was first investigated at age 17 for asymptomatic hyperCKaemia, with values of 3000-3500 UI/L. Muscle MRI of the lower limbs showed bilateral hypotrophy and early fatty changes of the *gastrocnemius medialis*, which was also hyperintense on T2-STIR sequences (Figure 1a). Muscle biopsy obtained from the same muscle displayed dystrophic changes with normal immunostaining for conventional sarcolemmal proteins (data not shown). Regarding the cardiological features, the patient did neither complain of suspicious symptoms (fatigue, dyspnoea, dizziness, palpitations) nor showed any structural or functional abnormality at baseline and follow-up visits. No arrhythmias were detected at baseline ECG and 24-hour Holter monitoring. The echocardiogram and the cardiac MRI showed a structurally normal heart. After almost four years of follow-up, he only complained about a vasovagal syncope triggered by emotional stress, while his Holter ECG showed a para-physiological sinus bradycardia with normal chronotropic competence during the day; the other clinical and instrumental cardiac features remained unchanged. The patient did not agree to undergo invasive tests such as electrophysiological study and loop recorder implantation, which were proposed.

Patient 2 (PT2) had a clinical onset at age 47 with myalgias and burning pain in the lower limbs. After one year, he underwent renal transplantation for chronic kidney failure, likely due to hypertensive nephropathy. One month after transplantation, he developed proximal lower limb weakness, which rapidly progressed in the following years. His CK level was above 6000 UI/L, with subsequent fluctuations between 2500 and 9000 UI/L, and based upon a suspicion of an immune-mediated myopathy, he was treated with steroids, together with one infusion of intravenous immunoglobulins and his chronic cyclosporine treatment, without benefit. Muscle imaging showed fatty replacement of *gluteus minimus, adductor longus*, and *magnus*, the left *semimembranosus* and *gastrocnemius medialis* bilaterally, together with relatively widespread abnormalities on T2-STIR images in the thigh muscles, especially in the anterior compartment, and in the *gastrocnemii*. An assay for myositis-specific antibodies turned out to be negative. At age 51, he could climb stairs using a handrail but could not raise from a chair without the use of arms. On physical examination, there was weakness of hip and knee flexion on the left side (Medical Research Council grade 4) and of knee extension bilaterally (grade 3). Electromyography was myopathic and nerve conduction studies were normal. Muscle biopsy from the right *vastus lateralis* showed, alongside myopathic changes, increased endomysial and perimysial fibrosis, and several necrotic fibres with myophagias; hypotrophic round fibres often concentrated in some fascicles; several nuclear clumps, and almost type II fibre uniformity on ATPase stainings were present. HLA class I staining was positive only in necrotic and regenerating fibres, and a mild reduction of sarcolemmal staining for caveolin-3 could be appreciated. Regarding the cardiological features, the patient did not complain of palpitations and did not have any syncope. His ECG was within normal range, only showing mild left axis deviation, and he did not show any arrhythmias on 24-hour ECG monitoring; his heart rate was normal throughout the recording. The echocardiogram showed mild, non-pathological interventricular septum hypertrophy (13 mm), which could be explained by hypertension, normal biventricular function, and no functional or structural abnormalities. The patient could not complete cardiac MRI because of claustrophobia. He agreed to undergo electrophysiological study and loop recorder implantation, which however has not yet been performed.

The POPDC1 p.V183F mutation affects a residue that is strongly conserved (Phylo IP100 score =7·844) and is present in all three vertebrate POPDC isoforms, and is also found in invertebrate POPDC proteins (Figure 1b). In the model of the Popeye domain of POPDC1, V183 is located in one of the β-strands (β4) and part of the jelly roll fold forming the roof of the cAMP binding Popeye domain (Figure 1c). V183 faces into the core of the Popeye domain in close proximity to the predicted cAMP binding pocket. Direct contact between cAMP and V183 is not predicted, although, while the substitution preserves the hydrophobic character at this position, it is unclear if there is any impact on cAMP binding due to steric effects (Figure 1d). The increased steric demand of phenylalanine may have a structural impact through clashes with other side chains in the β-folds (Figure 1e). However, modelling the V183F mutation using Missense3D ^32^ did not predict any major structural aberrations to the Popeye domain.

Skeletal muscle biopsy material from both patients and from age and sex-matched controls (CT1 and CT2) were sectioned and stained for either POPDC1 or POPDC2. Sections were also stained for SGCA to mark the sarcolemma of the fibres and served as a control for changes in the expression of POPDC isoforms (Figure 2a, b), as previously reported.^11,17^ The expression level of POPDC protein and SGCA was measured, with the median level of each control sample set to one, and the differences between the patients and controls analysed. An approximate 20% reduction (*p*<0·0001) in the median SGCA-normalised POPDC1 staining intensity in the sarcolemma of fibres of PT1 was observed (0·790, 95% CI: 0·744, 0·855; m=167) compared to CT1 (1·000, 95% CI: 0·982, 1·012; n=161). A similar reduction of around 25% (*p*<0.0001) was seen in PT2 (0·755, 95% CI: 0·731, 0·771; n=835) compared to CT2 (1·000, 95% CI: 0·972, 1·024; n=681) (Figure 2c, d). Meanwhile, the SGCA-normalised level of POPDC2 in the sarcolemma was reduced by 35% (*p*<0·0001) in PT1 (0·650, 95% CI: 0·619, 0·681; n=167) compared to CT1 (1·000, 95% CI: 0·951, 1·033; n=138), with a slightly milder reduction of 24% (*p*<0·0001) between PT2 (0·763, 95% CI: 0·715, 0·810; n=453) and CT2 (1·000, 95% CI: 0·938, 1·077; n=339) (Figure 2c, d). Mild reductions in the non-normalised expression levels of both isoforms at the sarcolemma were found (except for POPDC2 in PT2), while an increase in the cytoplasmic concentrations of POPDC2 was also observed in both PT1 and PT2 (Supplemental Figure 1a, b, d, e). These changes led to mild reductions in the enrichment of POPDC1 and POPDC2 at the sarcolemma membrane compared to the cytoplasm, which is representative of how effectively the POPDC proteins are localised at the sarcolemma (Supplemental Figure 1c, f). The changes in POPDC1 and POPDC2 expression were highly variable between individual fibres, with many fibres from the patients resembling control fibres with respect to POPDC protein expression while others showed major differences. Irregular and variable fibre sizes and morphologies were seen in both patients (Figure 2a and b). The median fibre cross-sectional areas were lower in both patients, with an 80% drop (*p*<0·0001) between CT1 (2968 μm^2^, 95% CI: 2843, 3101; n=296) and PT1 (628 μm^2^, 95% CI: 586, 680; n=327) and 63% reduction (*p*<0·0001) between CT2 (3269 μm^2^, 95% CI: 3190, 3380; n=1020) and PT2 (1215 μm^2^, 95% CI: 1151, 1322; n=1288) (Supplemental Figure 1g).

**Figure 2.**
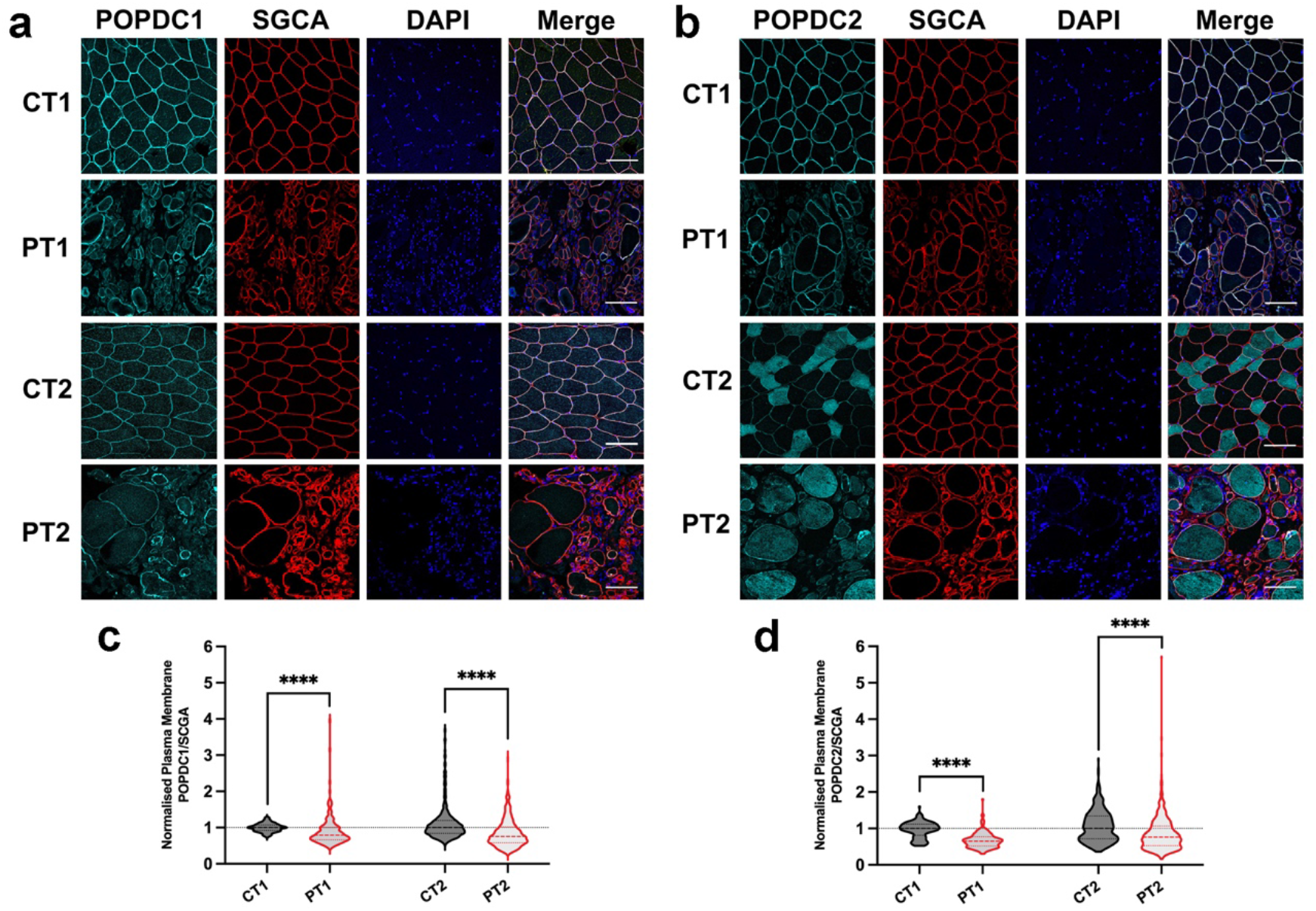
Membrane localisation of POPDC isoforms in muscle fibres expressing the POPDC1 p.V183F variant. (**a, b**) Transverse sections of skeletal muscle biopsies from PT1 and PT2 harbouring the *POPDC1* p.V183F variant and respective matched controls (CT1 and CT2) were stained for (**a**) POPDC1 or (**b**) PODPC2 along with SCGA serving as a sarcolemma marker. Scale bar: 100 μm. (**c, d**) The expression levels of (**c**) POPDC1 and (**d**) POPDC2 at the sarcolemma were normalised to SGCA and quantified in individual fibres. The number of sections (sec), images (img) and fibres (fib) analysed per group are as follows: CT1: POPDC1 - 2 sec, 5 img, 161 fib; POPDC2 - 2 sec, 5 img, 138 fib. PT1: POPDC1 - 1 sec, 4 img, 167 fib; POPDC2 - 1 sec, 4 img, 167 fibres. CT2: POPDC1 -1 sec, 13 img, 681 fib; POPDC2 - 2 sec, 8 img, 339 fib. PT2: POPDC1 - 3 sec, 20 img, 835 fib; POPDC2 - 3 sec, 14 img, 453 fib. The median POPDC/SGCA-level in each control biopsy was set to 1. Dashed lines indicate the normalised median and interquartile range. Data were analysed using Mann-Whitney test; **** *p*<0·0001.

### A POPDC1 p.Q153X mutation leads to a severe loss of POPDC1 and POPDC2 at the sarcolemma

A recently reported nonsense mutation in *BVES* (c.457>T, p.Q153X) is associated with early onset sinus bradycardia and atrioventricular block and high serum CK levels without clinical signs for LGMD.^20^ This mutation is predicted to lead to a truncation of POPDC1 within the Popeye domain, removing the cAMP binding domain and cytoplasmic C-terminal tail. A qualitative reduction in POPDC1 expression at the sarcolemma was reported in the affected patient, however no analysis of POPDC2 expression was performed.^20^ We have now quantified the changes in expression levels of POPDC1 and POPDC2 in the muscle fibres contained in biopsies from the index patient, along with a matched control (Figure 3a, b). The median SGCA-normalised POPDC1 intensity at the sarcolemma was around 75% lower (*p*<0·0001) in the patient (0·266, 95% CI: 0·250, 0·282; n=65) compared to the matched control (1·000, 95% CI: 0·962, 1·026; n=238). The normalised POPDC2 sarcolemmal level was even further reduced, by 88% (*p*<0·0001), between the patient (0·124, 95% CI: 0·116, 0·130; n=70) and the control (1·000, 95% CI: 0·970, 1·038; n=163). (Figure 3c, d). The absolute changes in POPDC1 and POPDC2 staining intensity at the sarcolemma were similar (Supplemental Fig 2a, d). A small decrease in POPDC1 and a moderate increase in POPDC2 were seen in the cytoplasmic levels (Supplemental Figure 2b, e). This led to a highly consistent reduction in the enrichment of POPDC1 and POPDC2 at the sarcolemma of the patient’s muscle fibres, with minimal variability (Supplemental Figure 2c, f). It was also noted that there was a greater than 2·5-fold increase (*p*<0·0001) in the cross-sectional area of muscle fibres between the control (2031 μm^2^, 95% CI: 1968, 2079, n=401) and patient biopsies (5283 μm^2^, 95% CI: 4707, 5952, n=135) (Supplemental Figure 2g).

**Figure 3.**
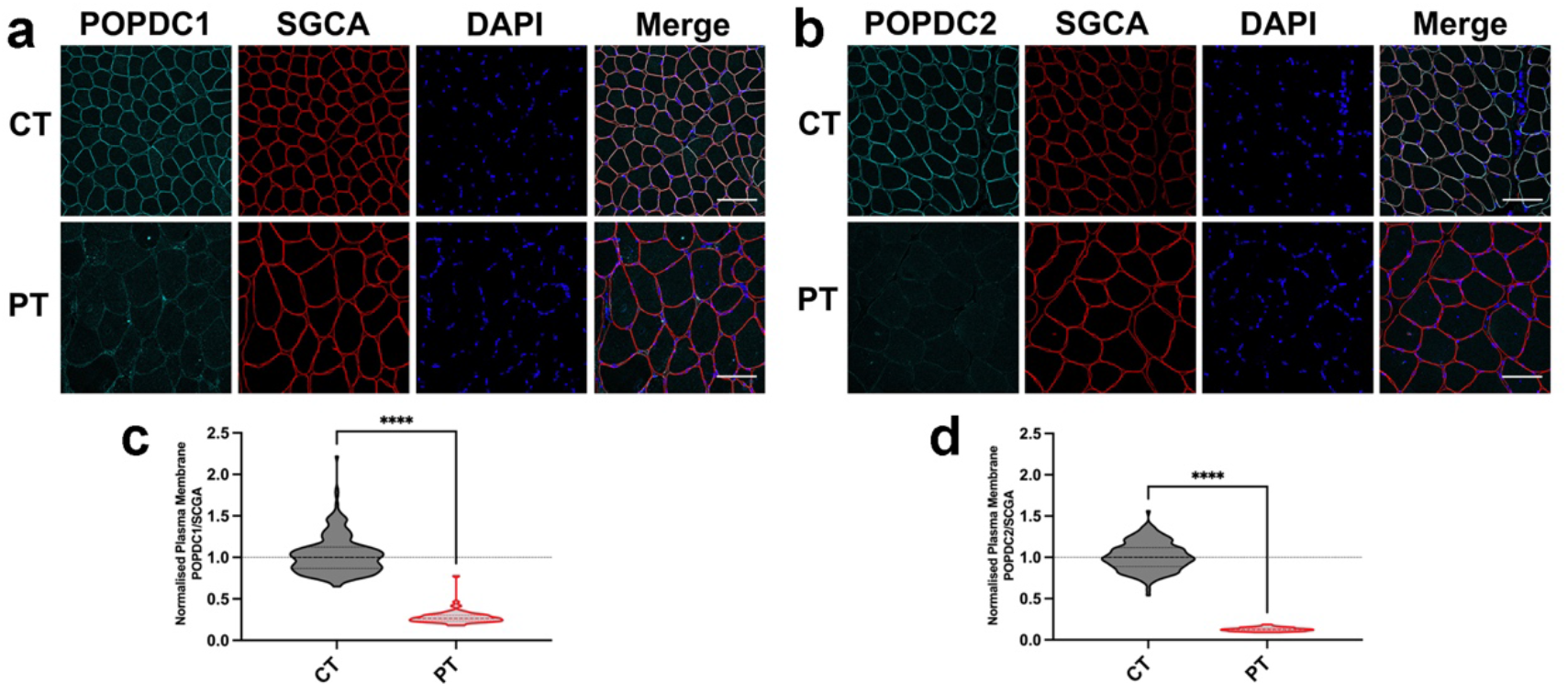
The expression of POPDC1 and POPDC2 is greatly reduced at the sarcolemma of skeletal muscle fibres expressing *POPDC1* p.Q153X. (**a, b**) Transverse sections of skeletal muscle biopsies from a patient (PT) carrying the *POPDC1* p.Q153X variant in homozygosity and a matched control (CT) were stained for (**a**) POPDC1 or (**b**) PODPC2, along with SCGA as a sarcolemma marker. Scale bar: 100 μm. (**c, d**) The expression levels of (**c**) POPDC1 and (**d**) POPDC2 in the sarcolemma normalised to SGCA, were quantified in individual fibres. The number of sections (sec), images (img) and fibres (fib) analysed per group are as follows: CT: POPDC1 - 1 sec, 4 img, 238 fib; POPDC2 - 1 sec, 4 img, 163 fib. PT: POPDC1 - 1 sec, 3 img, 65 fib; POPDC2 - 1 sec, 3 img, 70 fib. The median POPDC/SGCA-level in each control biopsy was set to 1. Dashed lines indicate the normalised median and interquartile range. Data were analysed using Mann-Whitney test; **** *p*<0·0001.

### Muscle fibres of *Popdc2*^*W188X/W188X*^ mutants show a loss of POPDC1 and POPDC2 at the sarcolemma

A heterozygous *POPDC2* (c.563G>A, p.W188X) mutation was previously reported in patients displaying atrioventricular block.^14^ No investigations into changes in the sarcolemmal expression of POPDC1 or the POPDC2 in skeletal muscle or heart tissue were performed due to a lack of biopsy material. We have created a homozygous *Popdc2*^W188X/W188X^ model mouse,^14^ from which the *gastrocnemius* of the mutant mouse and wild-type (WT) controls was taken and sections were stained for POPDC1 (Figure 4a) or POPDC2 (Figure 4b) along with SGCA, and the differences in staining compared. A 56% reduction (*p*<0.0001) was seen in the median SGCA-normalised POPDC1 intensity at the sarcolemma of each muscle fibre of the mutant (0·440, 95% CI: 0·416, 0·467, n=95) and WT (1·000, 95% CI: 0·939, 1·043; n=164) (Figure 4c). POPDC2 at the sarcolemma was reduced by 43% (*p*<0·0001) in the *Popdc2*^W188X/W188X^ mutant (0·471, 95% CI: 0·523, 0·661; n=93) and WT (1·000, 0·976, 1·050; n=143) (Figure 4d). The absolute change in POPDC1 and POPDC2 at the sarcolemma was similar (Supplemental Figure 3a, d), while only very mild changes in cytoplasmic levels of both POPDC isoforms were seen (Supplemental Figure 3b, e). This resulted in a lowering of the excess of POPDC1 and POPDC2 at the sarcolemma compared to the cytoplasm, with minimal variability (Supplemental Figure 3c, f). No major aberrations in fibre morphology were observed (Figure 4a, b), although a 10% reduction in the average fibre cross-sectional area (*p*=0·023) was seen in the mutant compared to wild type (Supplemental Figure 3g).

**Figure 4.**
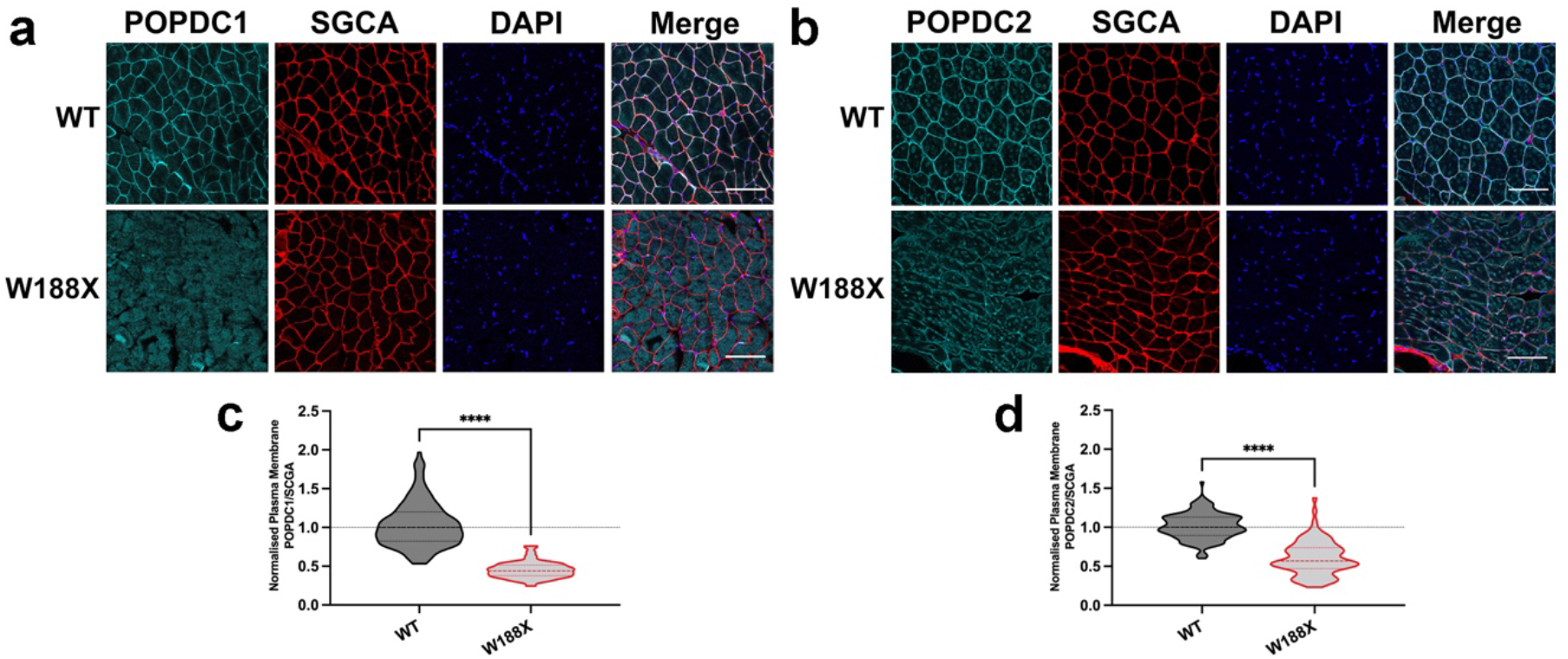
Intermediate reduction of POPDC1 and POPDC2 expression at the sarcolemma of skeletal muscle fibres of a homozygous *Popdc2* p.W188X knock-in mouse. (**a, b**) Transverse sections of the *m. gastrocnemius* of a 3-month-old homozygous *Popdc2* p.W188X knock-in mouse (W188X) and a wild-type control (WT) were stained for (**a**) POPDC1 or (**b**) PODPC2, along with SCGA as a sarcolemma marker. Scale bar: 100 μm. **(c, d**) The expression levels of (**c**) POPDC1 and (**d**) POPDC2 in the sarcolemma, normalised to SCGA, were quantified in individual fibres. The number of sections (sec), images (img) and fibres (fib) analysed per group are as follows: WT: POPDC1 - one sec, three img, 164 fib; POPDC2 - one sec, three img, 143 fib. W188X: POPDC1 - one sec, four img, 95 fib; POPDC2 - one sec, four img, 93 fib. The median POPDC/SGCA-level in each control biopsy was set to 1. Dashed lines indicate the normalised median and interquartile range. The control and homozygous mutant pairs were compared using a Mann-Whitney test; **** *p*<0·0001.

### Different POPDC mutations have a variable impact on sarcolemmal expression of POPDC1 and POPDC2

While all of the here studied mutations led to a reduction in the sarcolemmal expression level of POPDC1 and POPDC2, the effects were variable (Figure 5). Comparing the changes between each biopsy and its respective matched controls showed that the reduction in POPDC1 and POPDC2 levels at the sarcolemma in the case of the two patients carrying the POPDC1 p.V183F variant was significantly less than that seen in the patient possessing the POPDC1 p.Q153X mutation (*p*<0·0001; Figure 5). While there was no difference in the effect on POPDC1 across the two V183F patients (*p*=0·078; Figure 5), the reduction in POPDC2 was around 10% greater in PT1 (p=0·0030; Figure 5). The effect in the *Popdc2*^W188X/W188X^ mouse was less severe than in the patient expressing POPDC1 p.Q153X with respect to the loss of POPDC1 (*p*=0·016; Figure 5) and POPDC2 (*p*<0·0001; Figure 5). The effect on POPDC1 expression was however greater in the *Popdc2*^W188X/W188X^ mouse mutant than in both patients expressing POPDC1 p.V183F as well as for POPDC2 in case of PT2 carrying the POPDC1 p.V183F variant (*p*<0·0001; Figure 5). No difference was seen between the impact on POPDC2 sarcolemmal expression in case of PT1 carrying the POPDC1 p.V183F mutation and the *Popdc2*^W188X/W188X^ mouse mutant (*p*=0·32; Figure 5).

**Figure 5:**
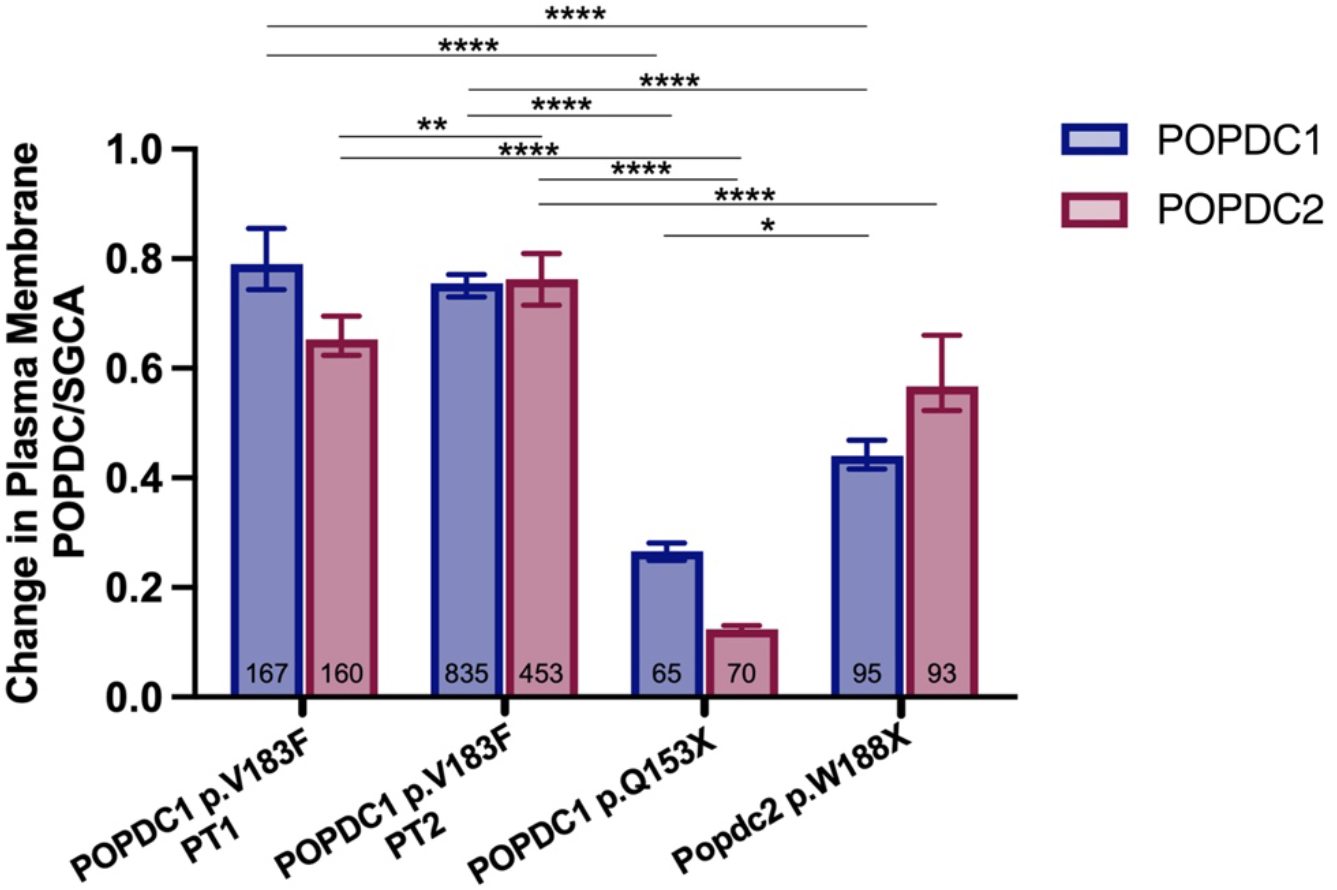
POPDC mutations have varying impacts on POPDC1 and POPDC2 sarcolemmal expression in skeletal muscle. Comparison of the fold change in POPDC1 and POPDC2 expression, normalised to SGCA, in the sarcolemma of skeletal muscle fibres from biopsy material of patients carrying the *POPDC1* p.V183F and *POPDC1* p.Q153X variants and a homozygous *Popdc2* p.W188X knock-in mouse compared to matched controls or wild-type mouse. The total number of fibres analysed are shown at the base of each bar. Data is displayed as median ± 95% CI. POPDC1 and POPDC2 values were compared using Kruskal-Wallis followed by Dunn’s multiple comparisons test; * *p*<0·05, ** *p*<0·01, **** *p*<0·0001.

### The plasma membrane expression and trafficking of POPDC1 and POPDC2 is dependent on each other

The above findings, as well of those from other previously reported patients,^11,17,19^ suggest that a mutation in POPDC1 can alter the subcellular expression pattern of POPDC2, and vice versa, in skeletal muscle fibres. To investigate if the membrane expression of POPDC1 and POPDC2 is indeed dependent on each other, HEK293 cells were transiently transfected with either POPDC1 or POPDC2 possessing C-terminal ECFP and EYFP tags, respectively (Figure 6a). Individual cells were segmented into cytoplasm and plasma membrane compartments using the lipophilic dye dioctadecyl-3,3,3,3-tetramethylindodicarbocyanine (DiD) to mark the plasma membrane and Hoechst-33342 to demarcate the nucleus. The extent of the plasma membrane localisation of each protein was quantified by determining the relative level of each protein at the plasma membrane compared to the cytoplasm. The median level of localisation of each protein across cells when singly expressed, or when co-expressed with the other POPDC isoform, was compared. When singly expressed, POPDC1 (0·565, 95% CI: 0·502, 0·713; n=15) and POPDC2 (0·506, 95% CI: 0·403, 0·773, n=15) were almost half as concentrated in the plasma membrane compared to the cytoplasm (Figure 6b). However, when co-expressed, POPDC1 (4·461, 95% CI: 3·692, 5·536; n=46) and POPDC2 (5·666, 95% CI: 4·254, 6·160, n=46) were both effectively localised to the plasma membrane at levels significantly above the single expression conditions (*p*<0·0001; Figure 6b). No significant difference in cytoplasmic levels of POPDC1 and POPDC2 between the two groups was seen (Supplemental Figure 4a). However, it was found that POPDC1 and POPDC2 plasma membrane expression when solely expressed was around 30% and 10% of the level observed in the co-expression system, respectively (*p*<0·0001; Supplemental Figure 4b).

**Figure 6.**
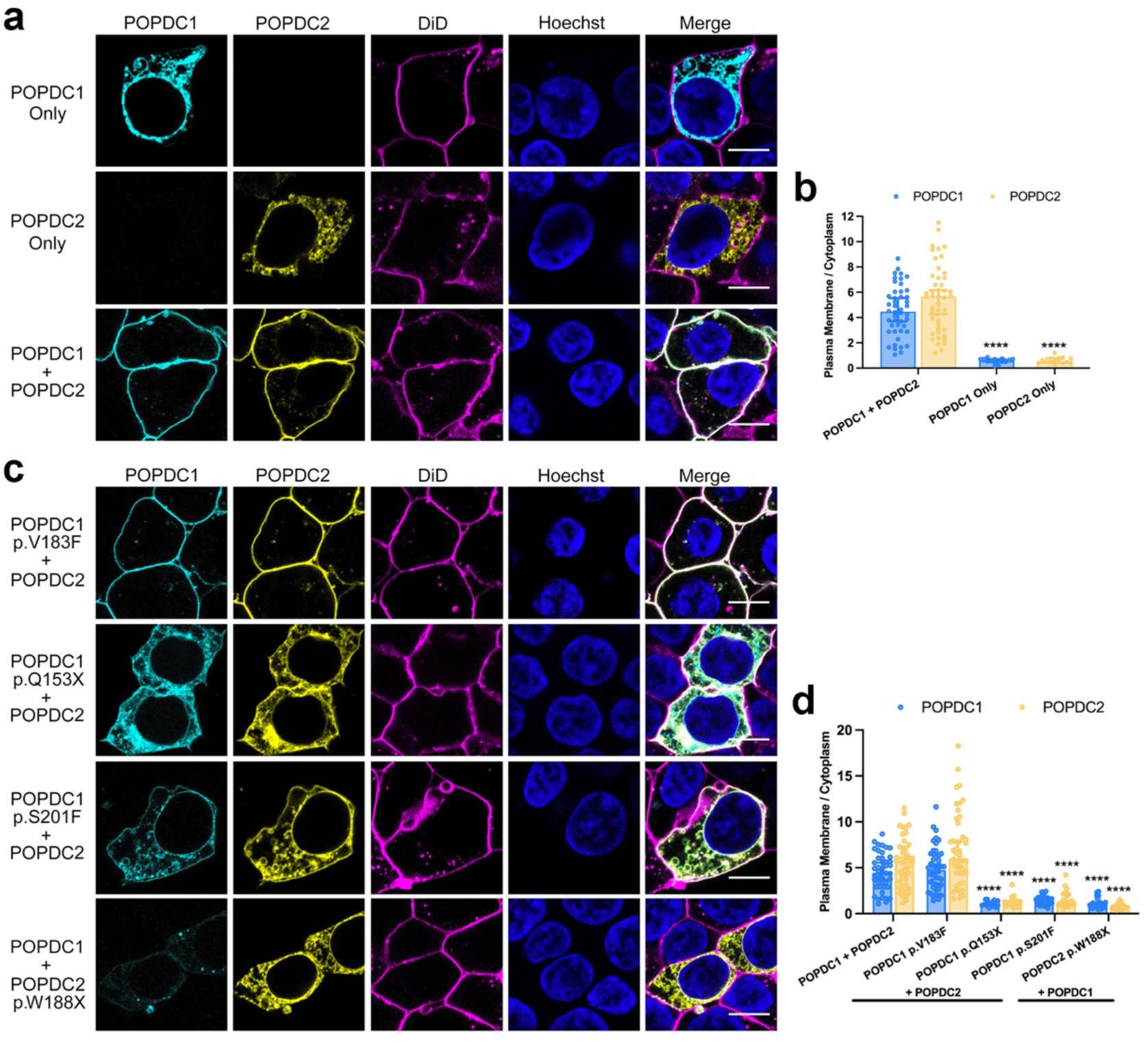
Co-expression of POPDC1 and POPDC2 is required for plasma membrane localisation in HEK293 cells. (**a**) POPDC1-ECFP, POPDC2-EYFP, or both were transiently expressed in HEK293 cells. The plasma membrane was marked using DiD. Scale bar: 10 μm. (**b**) Bar graph of the ratio of plasma membrane to cytoplasm expression of POPDC1 or POPDC2 (POPDC1-ECFP: n=15, POPDC2-EYFP: n=15, POPDC1-ECFP + POPDC2-EYP: n=46; min., N=2). Bars show median ± 95% CI. The groups were compared using a Mann-Whitney test; **** *p*<0·0001. (**c**) POPDC1 V183F-ECFP, Q153X-ECFP and S201F-ECFP and POPDC2 W188X-EYFP constructs were co-expressed with the appropriate wild-type POPDC partner in HEK293 cells. The plasma membrane was marked using DiD. Scale bar: 10 μm. (**d**) Bar graph of the ratio of plasma membrane to cytoplasm expression of POPDC1 or POPDC2 in the presence of the different POPDC1 and POPDC2 mutant proteins (WT n=46, V183F n=47, Q153X n=17, S201F n=22, W188X n=24, min., N=2). Identical data are shown in (**b**) and (**d**) for the expression levels after co-transfection of both wild-type constructs. Bars show median ± 95% CI. Groups were compared using Kruskal-Wallis followed by Dunn’s test using the wild-type pair for comparison; **** *p*<0·0001

The effect of a set of clinically identified POPDC mutations on the subcellular expression of POPDC1 and POPDC2 in HEK293 cells was then investigated. The POPDC1 p.V183F and p.Q153X mutations were introduced into the POPDC1-ECFP construct. Additionally, a POPDC1 p.S201F mutant was tested, having previously been reported to cause a significant loss in the plasma membrane expression of POPDC1 and POPDC2 in the muscle fibres of patients.^11^ The POPDC2 p.W188X mutation was introduced into the POPDC2-EYFP construct. Each mutant construct was co-transfected with its corresponding wild-type partner and the change in the median plasma membrane localisation of each protein, compared to the double wild-type expression, was determined. A significant reduction in the plasma membrane localisation of POPDC1 was seen in the presence of POPDC1 p.Q153X (0·967, 95% CI: 0·916, 1·243; n=17), POPDC1 p.S201F (1·520, 95% CI: 1·012, 1·735; n=22), and POPDC2 p.W188X (0·983, 95% CI: 0·830, 1·153; n=24) compared to the wild-type pair (all *p*<0·0001; Figure 6d). Likewise, significant drops in the plasma membrane enrichment of POPDC2 compared to the wild-type pair was seen for POPDC1 p.Q153X (1·239, 95% CI: 0·973, 1·369; n=17), p.S201F (1·202, 95% CI: 0·911, 2·078; n=22), and POPDC2 p.W188X (0·682, 95% CI: 0·573, 0·872; n=24) (all *p*<0·0001; Figure 6d). In contrast, the POPDC1 p.V183F mutation led to no significant changes in POPDC1 or POPDC2 plasma membrane localisation (Figure 6d). The POPDC1 p.Q153X mutation led to an increased accumulation of POPDC1 (*p*=0·015) and POPDC2 (*p*=0·042) in the cytoplasm, while POPDC2 p.W188X led to an increase in the intracellular localisation of the mutant POPDC2 protein (*p*<0·0001) (Supplemental Figure 4c).

### Direct interaction of POPDC1 and POPDC2

The above results led us to search for evidence of a direct interaction between POPDC1 and POPDC2. Both POPDC1 and POPDC2 are prominently expressed in cardiac and skeletal muscle ^1,2^ and immunostaining of isolated ventricular cardiac myocytes revealed overlapping expression domains for both isoforms (Figure 7a). To identify any interactions between POPDC1 and POPDC2 in their native environment, a proximity ligation assay (PLA) was carried out using sections from mouse atrium and ventricle from wild-type mice, with ventricular tissue of *Popdc1*^−/-^/*Popdc2*^−/-^ mutants serving as negative control (Figure 7b). PLA signals were observed in atrial and ventricular sections of wild-type hearts, whereas no signal was present in sections of *Popdc1*^−/-^/*Popdc2*^−/-^ mutants. While PLA signals were observed in cardiac myocytes of both chambers, the subcellular localisations differed between atrial and ventricular myocytes. In atrial myocytes, PLA signals were mostly localised at the sarcolemma, whereas in ventricular myocytes signals were found at the sarcolemma and within the cell boundaries. POPDC1 and POPDC2 have both been shown to reside in the sarcolemma and in the t-tubules of cardiomyocytes,^8^ with the higher level of t-tubules present in ventricular cardiomyocytes ^33^ likely contributing to the observed chamber-specific differences. This shows that POPDC1 and POPDC2 form complexes at the sarcolemma of cardiomyocytes. To confirm the interaction of POPDC1 and POPDC2, POPDC2 possessing a C-terminal FLAG tag was co-expressed with a POPDC1-Myc construct in COS-7 cells. In addition, POPDC2-FLAG was also co-expressed with POPDC3-Myc. Cell lysates were precipitated with a Myc-tag antibody and subjected to Western blot analysis using FLAG-tag antibody. POPDC1 was found to specifically co-precipitate with POPDC2, but this was not the case for POPDC3 (Figure 7c). Performing the same experiment with POPDC1 carrying a C-terminal HA-tag and POPDC3 with a FLAG-tag demonstrated that POPDC3 can be co-precipitated with POPDC1 (Figure 7d). This suggests that POPDC1 undergoes complex formation with POPDC2 and POPDC3, but no interaction is detectable between POPDC2 and POPDC3. The interaction of POPDC1 and POPDC2 was also further demonstrated to occur in native tissue by co-immunoprecipitation of POPDC1 from mouse heart lysates using a POPDC2 antibody (Figure 7e). The co-precipitation of POPDC1 did not occur when lysates were used from the hearts of *Popdc2* null mutant mice.

**Figure 7.**
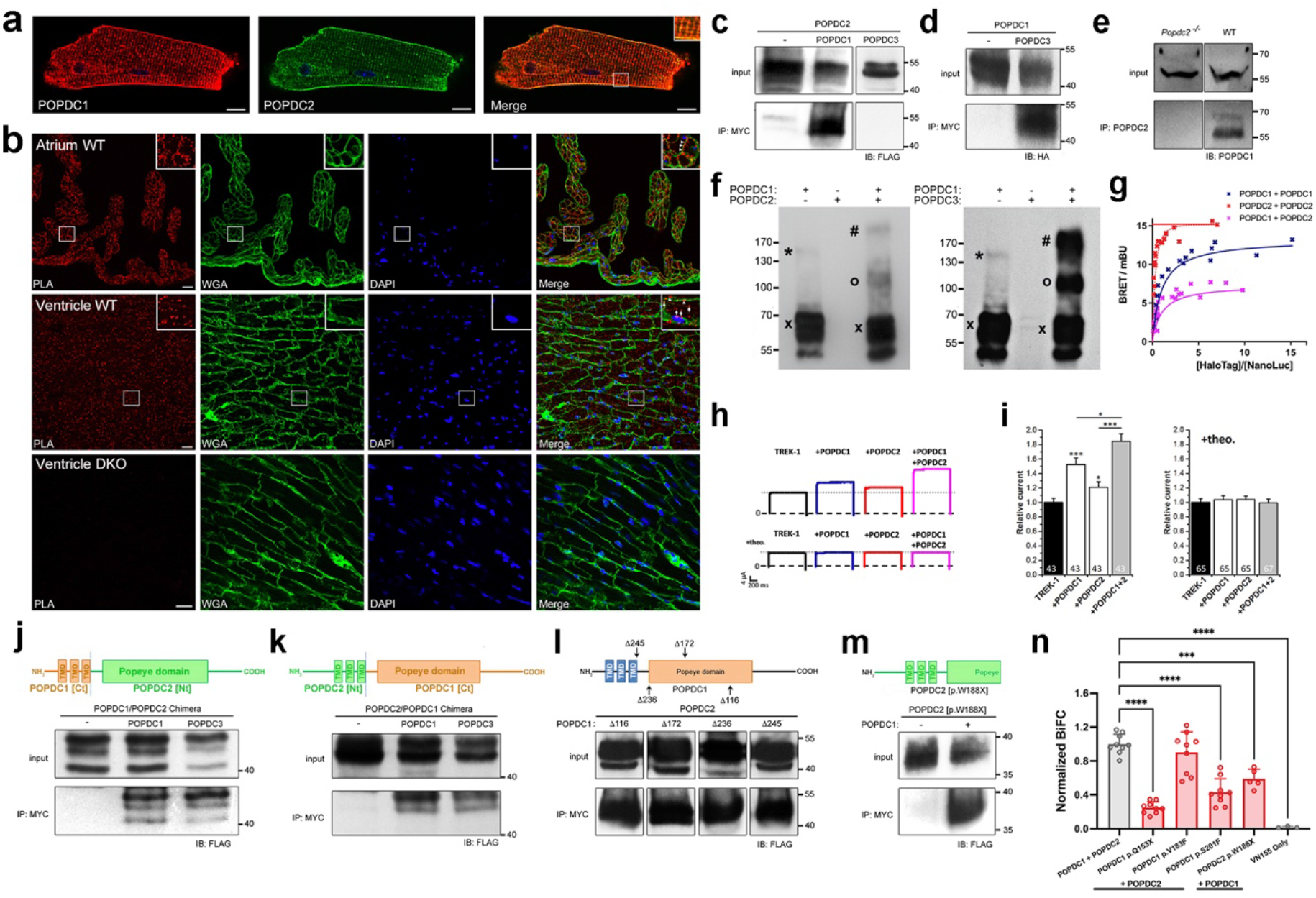
POPDC1 and POPDC2 undergo heteromeric complex formation. (**a**) Confocal microscopy of isolated mouse ventricular cardiomyocytes immunostained for POPDC1 (red) and POPDC2 (green). (**b**) Proximity ligation analysis (PLA) of POPDC1 and POPDC2 in transverse sections of right atrium and left ventricle isolated from wild-type and *Popdc1*/*Popdc2* double knockout mice. WGA staining (green label) was used to mark the sarcolemma and DAPI for the staining of nuclei. Red puncta are indicative of interactions between POPDC1 and POPDC2. Insets show magnifications of the indicated areas. (**c, d**) Co-precipitation analysis in COS-7 cells after co-transfection of (**c**) POPDC2-FLAG alone or together with POPDC1-Myc or POPDC3-Myc, respectively, or (**d**) POPDC1-HA alone or together with POPDC3-Myc. (**e**) Co-precipitation analysis of protein lysates of ventricular tissue isolated from *Popdc2* null mutant and wild-type mice. (**f**) Western blot analysis of lysates from COS-7 cells expressing POPDC1-CFP and/or POPDC2-FLAG (left blot), or POPDC1-CFP and/or POPDC3-FLAG (right blot). Bands were tentatively assigned as (x) monomer, (*) homodimer, (o) heterodimer or (#) heterotetramer based on their molecular weight. (**g**) Quantitative BRET saturation curves of POPDC1 and POPDC2 homo- and heteromeric complexes. (**h**) Examples of TREK-1 current in response to a voltage jump from −80 to 40 mV and (**i)** relative current amplitudes of *Xenopus laevis* oocytes expressing TREK-1 alone or together with POPDC1, POPDC2, or both isoforms together. Oocytes were incubated for 48-hrs in storage solution without or with theophylline (+theo). In (**i**) the number of oocytes for each experimental condition are given in each graph. Data are presented as mean ± SEM. * *p* <0·05, *** *p* < 0·001. (**j, k**) Chimeric constructs containing either (**j**) the N-terminal and transmembrane domains of POPDC1 (POPDC1 [Nt]) and the cytoplasmic region of POPDC2 (POPDC2[Ct]) or (**k**) the N-terminal and transmembrane domains of POPDC2 (POPDC2 [Nt]) and the cytoplasmic region of POPDC1 (POPDC1 [Ct]) were subjected to co-precipitation analysis after co-transfection together with POPDC1 or POPDC3 in COS-7 cells. (**l**) A series of truncations were introduced into POPDC1-Myc and subjected to co-precipitation analysis in COS-7 cells after co-transfection with POPDC2-FLAG. (**m**) A POPDC2-FLAG construct truncated to residue W188 was subjected to co-precipitation analysis in COS-7 cells after co-expression with POPDC1-Myc. (**n**) BiFC signal between wild-type POPDC1 and POPDC2, or in the presence of the POPDC1 p.V183F, p.Q153X, p.S201F and POPDC2 p.W188X mutations in HEK293 cells. POPDC1-VN155 + POPDC2-VN155 was used as a negative control. 5-9 images (approx. 50-100 cells per image) were taken per group across a min. of 2 transfections. Data is shown as mean ± SD. Groups were compared using One way ANOVA followed by Dunnett’s test using the wild-type pair as a comparison; *** *p*<0·001, **** *p*<0·0001.

When lysates of COS-7 cells expressing POPDC1 and POPDC2 were examined using SDS-PAGE followed by a Western blot, evidence of POPDC1 forming hetero-oligomers with POPDC2 was seen. Both POPDC1 and POPDC2 isoforms have very similar molecular weights (POPDC1: 41·5 kDa, POPDC2: 40·5 kDa). Therefore, POPDC1 was tagged at the C-terminal with CFP (29 kDa), while POPDC2 was fused with FLAG (1 kDa), to enable differentiation of homo- (expected: 140·5 kDa) and heterodimers (expected: 111·5 kDa) based on their molecular weight on the blot. Use of an anti-CFP antibody showed a differing pattern of bands when POPDC1 was expressed alone compared to co-expression with POPDC2. The majority of POPDC1 was in a monomeric state (multiple bands of approx. 70 kDa) in both groups, as would be expected given the presence of SDS. However, weaker bands corresponding to homodimers of POPDC1 (approx. 140 kDa), as well as those matching the expected molecular weight of POPDC1-POPDC2 heterodimers (approx. 110 kDa) and heterotetramers (approx. 220 kDa) were also seen, despite the denaturing conditions during electrophoresis (Figure 7f). Similar results were also seen when POPDC1 was co-expressed with POPDC3 (Figure 7f).

To investigate the stoichiometries of POPDC1-POPDC2 complexes within a cellular environment a type-1 quantitative BRET (qBRET) assay was employed, following the protocol reported by Felce et al.^25^ The NanoBRET platform ^34^ was utilised in the assay by tagging POPDC1 and POPDC2 at the C-terminus with NanoLuc luciferase (NL) or HaloTag (HT) fusion tags. These constructs were co-expressed in HEK293 cells at varying expression ratios, but at constant total expression levels (Supplemental Figure 5a-c) and the relationship to the BRET signal analysed. Firstly, POPDC1-NL and POPDC1-HT were co-transfected to try and identify if homomeric interactions were present as previously reported.^4,35^ A clear hyperbolic BRET saturation curve indicative of a dimer was apparent (Figure 7g). A rapidly saturating BRET curve was observed in the case of co-expression of POPDC2-NL and POPDC2-HT. Rapid saturation is a feature of higher order complexes, although such curve shapes make accurate determination of complex stoichiometry via type-1 qBRET studies difficult.^25^ When POPDC1-HT and POPDC2-NL were co-expressed, the BRET saturation curve produced fitted to a dimer model. This suggests that the major POPDC1-POPDC2 complex is a dimer. Such heterodimers would have to compete against the tendency of POPDC1, and likely POPDC2, to form homodimers, which suggests that the heteromeric-interaction of POPDC1 and POPDC2 is favoured.

We have previously shown that co-expression of POPDC1 or POPDC2 with the 2-pore domain potassium channel TREK-1 in *Xenopus laevis* oocytes leads to an increase in the outward K^+^ current compared to expression of TREK-1 alone.^5^ This was attributed to a direct, cAMP-sensitive interaction between POPDC proteins and TREK-1. Incubation of the cells with 8-Br-cAMP, or the phosphodiesterase inhibitor theophylline, abolishes the increase in current in the presence of POPDC1 or POPDC2, respectively.^5,11^ To test if the formation of heteromeric POPDC1-POPDC2 complexes could modulate TREK1 current, one or both POPDC isoforms were expressed in *Xenopus laevis* oocytes alongside TREK-1 and two electrode voltage-clamp measurements used to determine the outward K^+^ current from cells. As expected, co-expression of POPDC1 or POPDC2 with TREK-1 led to a significant increase in TREK-1 current compared to TREK-1 alone (Figure 7h, i). Furthermore, co-expression of POPDC1 and POPDC2 led to a significant additional increase in TREK-1 current above the levels observed with POPDC1 or POPDC2 alone. When the *Xenopus* oocytes were incubated with theophylline, the increase in TREK1 current after expression of POPDC1 and/or POPDC2 returned to baseline levels as previously reported.^5,11,14^

To help identify the domains responsible for the POPDC1-POPDC2 and POPDC1-POPDC3 interactions, two FLAG-tagged chimeras were constructed, consisting of the N-terminal and transmembrane domains of POPDC1 and the cytoplasmic region (including the Popeye domain) of POPDC2 and the inverse configuration (Figure 7j, k). Each chimera was co-expressed in COS-7 cells with Myc-tagged POPDC1 or POPDC3. Precipitation of cell lysates using Myc-antibody led to co-precipitation of both chimeras in all cases.

To further map the sites in POPDC1, which mediate the interaction between POPDC1 and POPDC2, co-immunoprecipitation experiments in COS-7 cells were repeated in the presence of various C-terminal truncations of POPDC1 (Figure 7l). The truncation mutants were C-terminally tagged with a Myc epitope and full-length POPDC2 with FLAG. Truncations were positioned to delete the C-terminal tail and end of the Popeye domain (Δ116), the C-terminal tail and half of the Popeye domain (Δ172), the C-terminal tail and the entire Popeye domain (Δ236) and the entire cytoplasmic region of POPDC1 (Δ245). After co-expression of these constructs with POPDC2, it was found that co-immunoprecipitation of POPDC2-FLAG was possible with all the truncation mutants, suggesting that the extracellular N-terminal region and transmembrane domains of POPDC1 were sufficient to form an interaction with POPDC2. A similar conclusion can probably be drawn for POPDC2, as the POPDC2 W188X mutant protein, which lacks the carboxy-terminal half of the Popeye domain and the carboxy terminus still retains the ability to interact with POPDC1 (Figure 7m). These results, and behaviour of the chimeric constructs, show that POPDC1-POPDC2 and POPDC1-POPDC3 interactions, occur at both the N-terminal/transmembrane domains and cytoplasmic portions of the proteins. It was shown above that the POPDC1 p.Q153X, p.S201F, and POPDC2 p.W188X mutations led to mislocalisation of POPDC1 and POPDC2 in HEK293 cells, as well as in skeletal muscle, while the POPDC1 p.V183F mutation had no effect in HEK293 cells and led to only mild changes in POPDC expression patterns in tissue. To investigate if a change in the interaction between POPDC1 and POPDC2 was responsible for this effect, a bimolecular fluorescence complementation (BiFC) assay was conducted (Figure 7n). POPDC1 and POPDC2 wild-type and mutant constructs were tagged at the C-terminal with the split Venus domains VC155 and VN155, respectively, and expressed in HEK293 cells. Interactions between VC155 and VN155 lead to reconstitution of the Venus fluorophore. POPDC1-VN155 expressed with POPDC2-VN155 was utilized as a negative control, while mRFP was co-transfected into the cells to act as an internal transfection control to which BiFC signals were normalised. The mRFP signal was also used to define areas containing transfected cells within confocal microscopy images (approx. 50-100 cells per image) from which the BiFC signal was measured (Supplemental Figure 6). As expected, wild-type POPDC1-VC155 and POPDC2-VC155 yielded a strong BiFC signal, which was set to equal 1, providing further evidence for the existence of POPDC1-POPDC2 complexes. No difference in the BiFC signal from POPDC1 p.V183F + POPDC2 compared to the wild-type pair was seen (n=9, *p*=0·54). However, a significant drop in the BiFC signal relative to the wild-type pair was observed in the presence of the POPDC1 p.Q153X (0·248, 95% CI: 0·195, 0·301; n=9), p.S201F (0·430, 95% CI: 0·307, 0·553; n=9), and POPDC2 p.W188X (0·590, 95% CI: 0·449, 0·730; n=5) variants (all *p*<0·0001). However, a detectable BiFC signal greatly above background was observed in all groups suggesting the POPDC1-POPDC2 interaction was not fully abolished.

### POPDC1 and POPDC2 may interact through a conserved interface in the αC-helix of the Popeye Domain

Having demonstrated that POPDC1 and POPDC2 interact through both their N-terminal/transmembrane and cytoplasmic regions, we focused on the role of the Popeye domain, which was previously reported to be involved in POPDC1 homomeric interactions.^4,35^ The cAMP binding domain of the prokaryotic cAMP-binding transcriptional regulator catabolite activator protein (CAP) shows the highest sequence similarity to the Popeye domain ^36^ and has therefore been used previously as a template for producing homology models of the Popeye domain.^5^ CAP protein monomers dimerize through an α-helix at the C-terminal end of their cyclic nucleotide binding domain (CNBD), known as the C-helix, via a set of hydrophobic residues ^37^ (Supplemental Figure 7a, b). As well as forming an interface between the CAP subunits, the C-helix also forms contacts with cAMP upon binding (Supplemental Figure 7c).^38,39^ The Popeye domains of POPDC1 and POPDC2 are predicted to be highly similar in structure to the CNBD of CAP, ^5^ and the protein structure of the CAP dimer (PDB: 1G6N^37^) was utilized as a template to model the POPDC1-POPDC2 heteromeric complex (Figure 8a). An α-helix is predicted to form at the C-terminal end of the Popeye domain and is referred to as the αC-helix in reference to the structures of PKA and other cAMP effector proteins.^40,41^ The model Popeye domain dimer possesses an interface between each αC-helix, analogous to the C-helix interface in CAP. Alignment of the amino acid sequences of the αC-helices of vertebrate POPDC1, POPDC2 and POPDC3, as well as the C-helix of CAP, reveals a high level of sequence conservation and in particular the invariant presence of a series of hydrophobic residues in each POPDC isoform, which, with the exception of one residue, were also present in the C-helix of CAP (Figure 8b). Some of these residues are known to be involved in CAP dimerization (Supplemental Figure 7b).^37,38^ These hydrophobic residues show very strong structural alignment across the predicted structures of the αC-helix in POPDC1, POPDC2, and POPDC3 (Figure 8c, d). The results shown above suggest that if normal POPDC1-POPDC2 interactions are disrupted, or absent, then the subcellular expression of both isoforms is altered. To determine if the CAP-aligned, highly conserved hydrophobic residues within the αC-helices of POPDC1 and POPDC2 are involved in complex formation, POPDC1-ECFP and POPDC2-EYFP were co-expressed in HEK293 cells, with each of the conserved hydrophobic residues in the αC-helix sequentially substituted to aspartic acid (Supplemental Figures 8a, 9a). These substitutions were designed to disrupt the hydrophobicity of the putative helix-helix interface through the introduction of a negative charge. The median plasma membrane localisation level of both POPDC isoforms across the cells was then determined as before and the difference to the wild-type pair analysed. It was found that POPDC1 mutations F249D (0·812, 95% CI: 0·584, 0·990; n=56), I253D (0·841, 95% CI: 0·571, 1·185; n=27) and I257D (0·625, 95% CI: 0·441, 0·741; n=40) led to severe mislocalisation of POPDC1 compared to the wild-type pair (3·676, 95% CI: 3·176, 4·504; n=34) (all *p*<0·0001, Figure 8e). These mutations had a similar effect on POPDC2 with F249D (0·845, 95% CI: 0·650, 0·982; n=56), I253D (1·202, 95% CI: 0·790, 2·060; n=27), and I257D (0·635, 95% CI: 0·488, 0·779; n=40) (all *p*<0·0001) all leading to a reduction in the plasma membrane localisation of POPDC2 compared to when co-expressed with wild-type POPDC1 (4·443, 95% CI: 3·902, 6·662; n=34). The other mutations within the αC-helix of POPDC1 did not lead to any significant changes in POPDC1 or POPDC2 plasma membrane localisation (Figure 8e). In POPDC2, the F233D (0·922, 95% CI: 0·743, 1·277; n=20), L237D (0·967, 95% CI: 0·750, 1·267; n=45), and I241D (0·735, 95% CI: 0·642, 0·922; n=42) mutations led to a significant disruption of POPDC1 plasma localization (all *p*<0·0001) compared to when expressed with wild-type POPDC2 (Figure 8f). These mutations also led to mislocalisation of POPDC2 itself: F233D (0·778, 95% CI: 0·651, 0·980; n=20), L237D (1·121, 95% CI: 0·860, 1·644; n=45), and I241D (1·146, 95% CI: 0·946, 1·479; n=42) (all *p*<0·0001). A mild reduction in POPDC2 plasma membrane localisation was also seen with the I229D mutation (2·828, 95% CI: 2·098, 3·337; n=48, *p*=0·049) (Figure 8f). The L245D, L261D, and L264D mutations in POPDC1 and the I229D, L245D, and L248D in POPDC2, had no effect on the subcellular expression of either protein (except the minor change in POPDC2 expression in case of I229D). A loss in absolute plasma membrane expression was commonly seen in mutations that led to mislocalisation of the proteins (Supplemental Figure 8b, 9b), with lesser or no changes in cytoplasmic levels observed (Supplemental Figure 8c, 9c). Residues that are aligned between POPDC1 and POPDC2 had very similar impacts on the plasma membrane localisation of both isoforms (Figure 8g, h). Mutation of the three residues at the core of the proposed αC-helices led to major losses in POPDC1 and POPDC2 plasma membrane localisation, while substitution of the residues at the N- and C-termini of the helices had no, or only minimal effect (Figure 8i).

**Figure 8.**
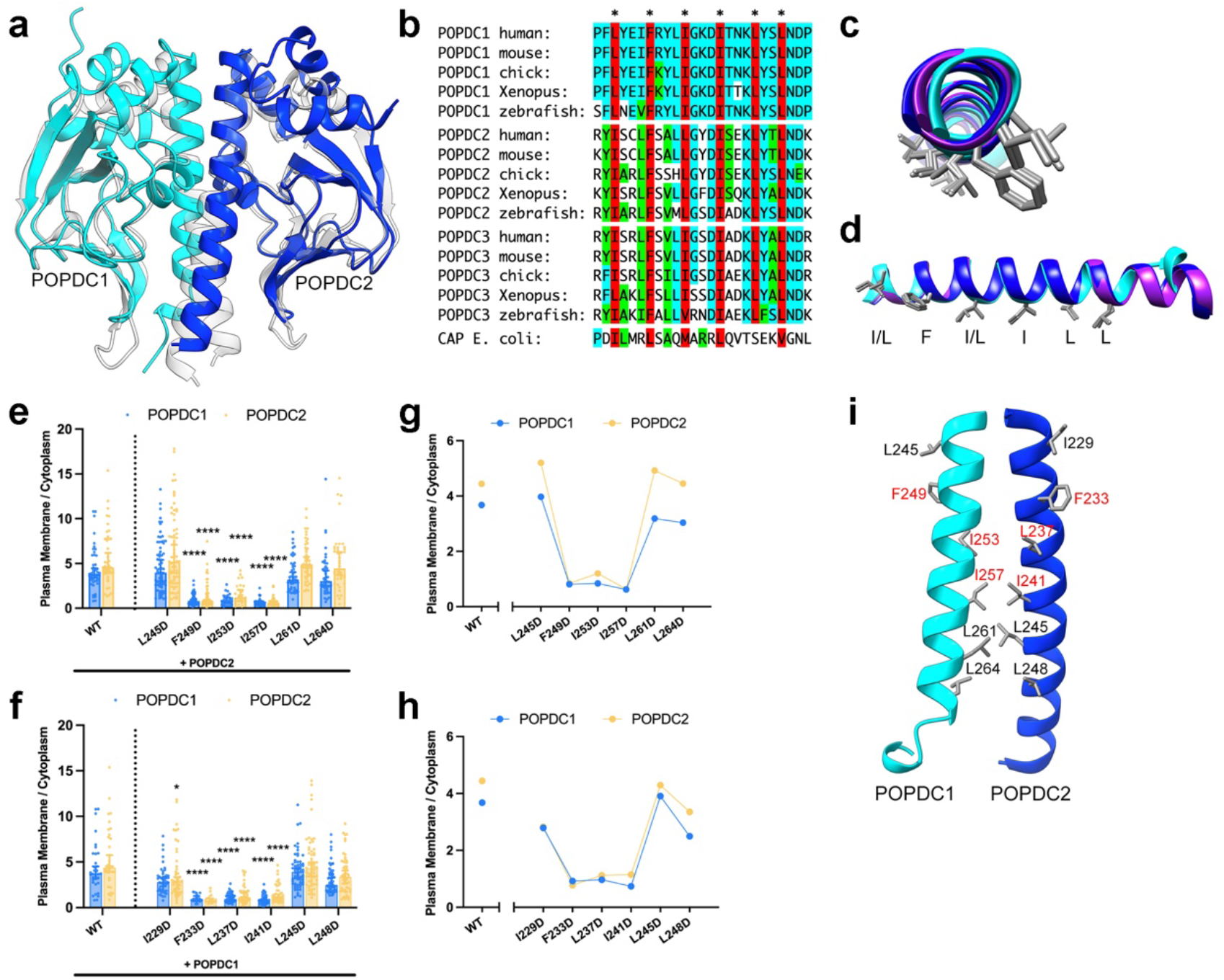
The αC-helix of the Popeye domain mediates heteromeric complex formation between POPDC1 and POPDC2. (**a**) A model of heteromeric complex formation of the Popeye domains of POPDC1 (cyan) and POPDC2 (blue) using the structure of a CAP dimer (translucent; PDB: 1G6N) as a template. (**b**) Sequence alignment of the αC-helix of POPDC1, POPDC2 and POPDC3 from multiple species and CAP. A set of highly conserved hydrophobic amino acids are highlighted in red. (**c**) Amino terminal view and (**d**) side on view of an overlay of the predicted αC-helical structures of POPDC1 (cyan), POPDC2 (blue) and POPDC3 (purple) with the side chains of the highly conserved hydrophobic residues shown. (**e, f**) The ratio of plasma membrane to cytoplasm expression levels of POPDC1-ECFP and POPDC2-EYFP in HEK293 cells, where either (**e**) POPDC1-ECFP or (**f**) POPDC2-EYFP was subjected to site-directed mutagenesis to introduce an aspartic acid in place of a conserved hydrophobic residue within the αC-helix sequence. Total number of cells analysed: POPDC1: L245D n = 71, F249D n = 56, I253D n = 27, I257D n = 40, L261D n = 47, L264D n = 42; POPDC2: I229D n = 48, F233D n = 20, L237D n = 45, I241D n = 42, L245D n = 63, L248D n = 56. Min. 2 transfections per group. Bars show median ± 95% CI. Groups were compared using Kruskal-Wallis followed by Dunn’s test using the wild-type pair as a comparison; ** *p* < 0·01, **** *p* < 0·0001. (**g, h**) The relationship between plasma membrane versus cytoplasm expression and each mutation in (**g**) POPDC1 and (**h**) POPDC2. (**i**) The predicted αC-helical Popeye domain interface between POPDC1 and POPDC2. Hydrophobic residues whose mutation to aspartic acid led to severely impaired plasma membrane localisation of both POPDC proteins are labelled in red.

## Discussion

The two probands homozygous for the p.V183F variant displayed a peculiar, skeletal muscle-restricted phenotype, largely different from what has been described in other patients carrying *BVES* variants,^11,17,18^ especially since cardiological workup has not yet revealed signs of heart disease in both cases. The phenotypes in these patients likely represent a milder end of the clinical spectrum associated with this gene, with this missense variant seeming to cause a disease starting in the posterior lower leg, later progressing to the proximal lower limbs, with relative sparing of the upper body. Muscle pain with high CK and distal onset in the posterior lower leg has been described in Japanese patients with biallelic nonsense *BVES* variants.^19^ Significant hyperCKaemia associated with muscle necrosis on biopsy, onset as a calf myopathy with subsequent fast progression in adulthood, and the distribution of MRI changes (*adductor longus* and *magnus, gluteus minimus* and *semimembranosus*, together with abnormalities on T2-STIR images and some degree of asymmetry) closely resembled LGMDR12.^42^ It is interesting in this regard that POPDC1 has been shown to physically interact with anoctamin 5.^43^ Recently, one additional patient with biallelic nonsense *BVES* variants, without signs of relevant arrhythmogenic heart disease at age 33 and with diffuse, asymmetric changes on T2-STIR MRI images, has been reported.^21^ Notably, in both our patients extensive genetic testing did not reveal concomitant potentially pathogenic variants in other genes causative for myopathies, and a superimposed inflammatory myopathy in PT2 was not supported by muscle pathology, response to treatment or serological data.

In several previously reported cases of patients carrying a variant in *BVES* and developing LGMDR25, a loss in sarcolemmal localisation of both POPDC1 and POPDC2 in skeletal muscle fibres was reported.^11,17,19^ We therefore investigated the effect of the *BVES* p.V183F variant on the sarcolemmal expression of POPDC1 and POPDC2 in these patients. The mild reductions in the sarcolemmal expression level for both isoforms we observed were in contrast to the skeletal muscle fibres of the patient carrying a homozygous *BVES* p.Q153X variant,^20^ which displayed a substantial loss of both isoforms at the sarcolemma resembling other investigated *BVES* variants.^11,17,19^ The homozygous *Popdc2*^W188X^ mouse carrying a mutation which has been recently identified in two families suffering from AV-block but having normal skeletal muscle,^14^ showed a reduction in sarcolemma localisation at a level intermediate to those seen for patients carrying *BVES* p.V183F and p.Q153X variants. Our results demonstrate that mutations in either POPDC1 or POPDC2 may result in impaired membrane localisation, suggesting that membrane trafficking is dependent on simultaneous expression of both proteins, but the severity of this effect is mutation-specific.

Previously, it has been reported that forced expression of POPDC1 in cell lines resulted mainly in intracellular expression with only a small amount of the transfected protein reaching the plasma membrane, which contrasts with native tissue where the protein is mainly localised at the sarcolemma.^4,11,44,45^ Our finding that co-expressing POPDC1 and POPDC2 in HEK293 cells yielded a high degree of plasma membrane localisation of both proteins support the notion that co-expression of POPDC1 and POPDC2 is required for proper plasma membrane transport. It also permitted the use of HEK293 cells as a model system for investigating POPDC protein subcellular expression patterns. The disruption of plasma membrane expression and localisation of POPDC1 and POPDC2 in HEK293 cells in the presence of POPDC1 p.Q153X, p.S201F, and POPDC2 p.W188X mutations, replicated their effects in native tissue.^11^ This suggests that the changes in subcellular localisation of POPDC proteins seen in patients carrying *BVES* or *POPDC2* variants are due to a fundamental effect on POPDC proteins and is not a tissue-specific response only present in striated muscle tissue.

We explain the requirement for POPDC1 and POPDC2 to be both present for normal plasma membrane localisation by the formation of heteromeric complexes, probably during translation or shortly thereafter. Direct POPDC1-POPDC2 interactions were shown to occur after forced expression as demonstrated by co-immunoprecipitation, qBRET and BiFC, with their presence confirmed in native tissue by PLA and co-immunoprecipitation in cardiac tissue. The qBRET experiments showed a preference for heteromeric POPDC1-POPDC2 dimers, while Western blot experiments suggested that higher order heteromers may be able to form. However, these assays utilise non-native overexpression systems, as well as denaturing conditions in the case of the Western blot, which may lead to non-native oligomer formation. Further investigations into POPDC complex stoichiometries using native tissue would be beneficial. Alongside disrupting the plasma membrane localisation of POPDC1 and POPDC2, the BiFC assay also showed that the POPDC1 p.Q153X, p.S201F and POPDC2 p.W188X mutations impaired the interaction between POPDC isoforms. The POPDC1 p.V183F mutation preserved plasma membrane expression of both POPDC1 and POPDC2 isoforms in HEK293 cells and did not affect their interaction according to the BiFC results. This provides evidence that disruption of the POPDC1-POPDC2 complex is responsible for the major changes in subcellular localisation in muscle tissue of patients.

The preservation of coimmunoprecipitation of POPDC1 and POPDC2 when POPDC1 was subjected to a series of truncations demonstrates that interactions occur through the N-terminal and/or transmembrane regions. Although the nature of the responsible interfaces at these sites is unknown, they likely involve hydrophobic interactions between the transmembrane domains (Supplemental Figure 10a). It is however clear from our analysis that this interface(s) are not sufficient for triggering membrane transport. The use of the *E. coli* CAP protein as a template for modelling the possible interaction of the Popeye domains of POPDC1 and POPDC2 suggested that the αC-helix within the Popeye domain of each protein may form an heteromeric interface. The set of highly conserved hydrophobic residues, and the modelled structures of the αC-helix, means that such an interface would likely be pseudo-symmetrical supporting its function as an interaction domain.^48^ Substitution of these hydrophobic residues with aspartic acid had specific and symmetrical effects on both proteins with residues F249, I253, and I257 in POPDC1, and F233, L237, and I241 in POPDC2, which are aligned with each other, shown to be particularly important for normal subcellular localisation in HEK293 cells. The αC-helix is lost in the case of the POPDC1 p.Q153X and POPDC2 p.W188X nonsense mutations, which would result in a total loss of αC-helix mediated interactions. In contrast, the missense POPDC1 p.V183F variant is predicted to be positioned distal to the αC-helix, which may explain its relatively mild effect on POPDC protein localisation.

The POPDC1 p.S201F missense mutation caused pronounced changes in plasma membrane localisation of POPDC1 and POPDC2 and disruption of POPDC1-POPDC2 complex formation despite being positioned outside the αC-helix. In CAP, the C-helix mediates both dimerization and cAMP binding, which leads to stabilisation of the helix.^37,38^ It is known that the S201F mutation reduces the cAMP binding affinity of POPDC1 by around 50%.^11^ Modelling of cAMP binding to the Popeye domain suggests the αC-helix may be involved in cAMP binding, particularly the highly conserved K260 and L264 residues (Supplemental Figure 10b). We also note that in the non-cAMP bound model of POPDC1, the side chains of K260 and D256, both within the αC-helix, are predicted to form a H-bond network with the hydroxyl group of S201, as well as the backbone of other residues within the phosphate binding cassette (PBC) (Supplemental Figure 10c), which may mediate αC-helix conformation. Therefore, cAMP binding may have an allosteric effect on POPDC protein interactions via the αC-helix, perhaps explaining the changes in subcellular localisation caused by the POPDC1 p.S201F variant. We note that the effect of the POPDC1/POPDC2 heteromeric complex on TREK-1 current was strongly influenced by raising cAMP levels through theophylline in *Xenopus* oocytes.^5^

While we still do not fully understand the mechanism underlying the parallel changes in the subcellular localisation of POPDC1 and POPDC2 in response to a single mutation, there is a strong precedent for the requirement of heteromeric interactions for correct trafficking of transmembrane proteins. Examples include the T cell antigen receptors^49^ and the GABA_B_ receptors,^50–54^ which require complete heteromeric interactions between the receptor complex subunits to avoid endoplasmatic reticulum (ER) retention and subsequent degradation. Several possible ER retention motifs are present within the Popeye domain and C-terminal domain of POPDC1 and POPDC2. Heteromeric Popeye domain interactions through the αC-helices may be required to mask ER retention motifs in POPDC1 and POPDC2 and permit correct movement of the complex to the plasma membrane. Loss of mutant protein may lead to reduced POPDC1-POPDC2 complex formation. A reduction of POPDC1 protein levels in the patient possessing the *BVES* (c.457C>T, p.Q153X) mutation was previously suggested to be due to nonsense mediated decay (NMD),^20^ although Rinné *et al*. detected *POPDC2* (c.563G>A, p.W188X) coding mRNA and POPDC2 protein in patient leukocytes, suggesting that NMD did not totally prevent POPDC2 expression in that case.^14^

It is likely that POPDC1-POPDC2 interactions are not totally abolished in the case of the investigated mutations, with the N-terminal/transmembrane interface less affected by the mutations within the Popeye domain, although the nature of the complex is likely altered. This is supported by the presence of BiFC signals which were above background in all cases. In native tissue, other factors likely influence POPDC protein expression and localisation; indeed in all cases investigated above, the mutations did not completely abolish POPDC expression at the plasma membrane of muscle fibres or HEK293 cells. While preservation of a POPDC1-POPDC2 interface outside the Popeye domain may prevent complete loss of both proteins at the membrane, interactions with other proteins may also be relevant. POPDC1 and POPDC2 show varying expression profiles across tissues and so are unlikely to always be present in stoichiometric amounts.^55^ It appears that POPDC proteins may form both homo- and hetero-oligomers, meaning there may be an equilibrium between these states within cells, with homomeric POPDC interactions previously suggested to be involved in mediating subcellular localisation.^56^ Interestingly, the homomeric interaction of POPDC1 was previously suggested to involve sequences immediately C-terminal to the αC-helix,^4,35^ suggesting that this interface may also support homomeric interactions. However, based on the results of the qBRET experiments, POPDC1 and POPDC2 seem to preferentially undergo heterodimer formation. We also identified an interaction between POPDC1 and POPDC3. This observation is of significance as POPDC1 and POPDC2 are co-expressed in heart and skeletal muscle, whereas in brain POPDC2 is absent but both POPDC1 and POPDC3 are co-expressed.^46,47^ Although we have not further investigated whether POPDC1 and POPDC3 expression would also support membrane localisation the high level of sequence homology, particularly of the αC-helix, between the isoforms suggests this may be possible.

Why POPDC hetero-oligomer formation is required is unclear but may relate to isoform-specific protein-protein interaction partners, meaning that POPDC heterodimer formation would permit assembly of particular complexes. Other possibilities include changes to cAMP binding affinity in the hetero-oligomer, which may be important for the proper function of POPDC proteins in the context of cAMP signalling. The spatial arrangement of POPDC proteins, which we have shown to be dependent on POPDC1-POPDC2 complex formation and sensitive to mutations in either protein, may influence the assembly of cAMP signalling complexes and nanodomains, which are vital in cAMP dependent signalling.^58,59^ Indeed, POPDC1 has been shown to form functionally important interactions with cAMP pathway proteins such as PDE4, AC9, and TREK-1 in the heart, the disruption of which may impact calcium transients, β-adrenergic signalling, and ion channel function.^12,13^ Such an effect could explain the cardiac arrhythmias which have been observed in many patients carrying *BVES* or *POPDC2* variants^11,14,17–21^ and may also be responsible for the muscular dystrophy phenotype associated with POPDC mutations, although the link to cAMP signalling has not yet been established in the case of skeletal muscle. However, POPDC1 and POPDC2 have been reported to interact with a number of proteins linked to sarcolemmal stability or repair such as dystrophin,^11^ dysferlin,^11^ Xin-related protein 1,^45^ annexin A5,^45^ anoctamin 5,^43^ and caveolin-3.^8^ Disruption of the PODPC1-POPDC2 complex may lead to altered function of these interaction partners and so contribute to the observed muscular pathologies and hyperCKaemia observed in patients.

Both homozygous *BVES* p.V183F patients reported here display only a muscular phenotype with no evidence for a cardiac arrhythmia or any other pathology in the heart, suggesting a tissue-specific effect. The V183F mutation affects a residue located in the β4-strand of the Popeye domain. Recently, a binding site for PDE4 was mapped to the β3-strand of POPDC1.^12^ By analogy, we hypothesize that the V183F mutation may potentially affect a binding site for an unknown interaction partner, which is essential for skeletal muscle but dispensable for the heart. Given the orientation of V183 into the core of the Popeye domain this would require the mutation to cause allosteric alterations to the protein. Such an effect would be in contrast to changes in cAMP affinity, as reported for the S201F mutation,^11^ which would be present in all tissues. In addition, the preservation of POPDC1-POPDC2 complex formation may explain the apparent normal POPDC1-POPDC2 function in the heart and the absence of a cardiac phenotype in these patients. Given the likely relevance of POPDC1-POPDC2 interactions for sarcolemmal expression, and impaired membrane localisation in patients carrying POPDC mutations, we recommend establishing if novel or previously reported mutations lead to changes in the interaction and localisation of the proteins in tissue biopsy and/or HEK293 cells. Unravelling the effects of a loss of POPDC1-POPDC2 complexes at the sarcolemma, particularly the changes in the array of POPDC-interacting proteins known to be required for normal skeletal muscle and cardiomyocyte function, and cAMP signalling, will be needed to understand the range of phenotypes seen in patients carrying POPDC gene mutations.

## Contributors

BU, GT, MP, ND, and TB designed parts of the study and supervised the experimental work. AHS, FB, IM, MS, RFRS, and SR performed all the experiments. AH, AS, GT, FP, and MS provided clinical data and expertise. AHS, GT, MS, and TB analysed the data and wrote the manuscript. ACS, BU, KC, ND, MP, and TB have verified the underlying data. All authors had access to the data and had reviewed the manuscript and approved the submitted version.

## Data sharing statement

The datasets generated during the current study will be made available from the corresponding author on reasonable request.

## Acknowledgement

Expert technical assistance of Mrs. Ursula Herbort-Brand is hereby gratefully acknowledged. We thank Stephen Rothery of the Facility for Imaging by Light Microscopy, Department of Medicine, Imperial College London who is in part funded by the British Heart Foundation (RE/18/4/34215) for his assistance and expertise in obtaining the sample images included in this work. This work was funded by the British Heart Foundation (PG/14/46/30911) to TB and the Deutsche Forschungsgemeinschaft (DE1482/9-1) to ND. AHS was funded by an EPSRC/British Heart Foundation co-funded Imperial Institute of Chemical Biology (ICB) Centre for Doctoral Training (CDT) PhD studentship (EP/S023518/1).

