## Supplemental Data for "Differential Effects of Mutations of Popeye Domain Containing Proteins on Heteromeric Interaction and Membrane Trafficking"

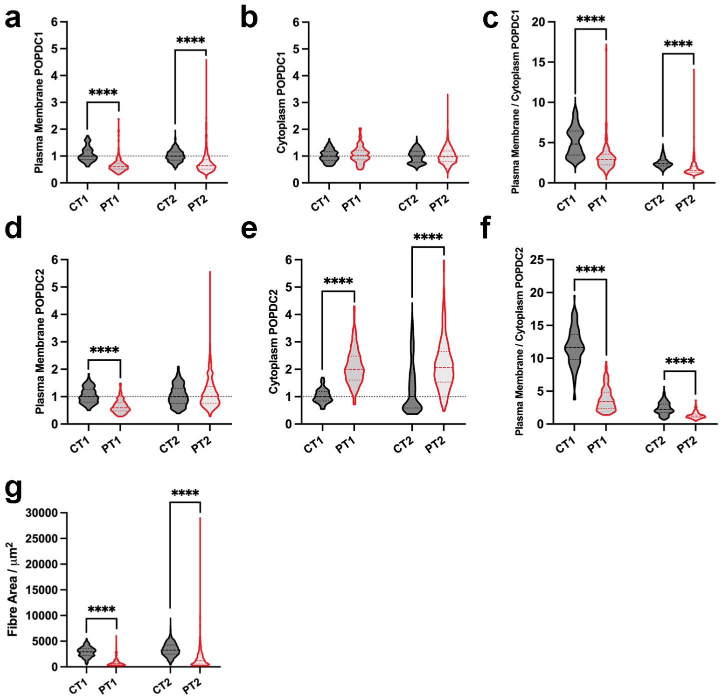


**Supplemental Figure 1. Expression analysis of POPDC1 and POPDC2 in skeletal muscle biopsies of patients carrying a *POPDC1* p.V183F mutation.** (**a**-**c**) POPDC1 and (**d**-**f**) POPDC2 expression in (**a**, **d**) sarcolemma, (**b**, **e**) cytoplasm and (**c**, **f**) the ratio of sarcolemma/cytoplasm expression as determined by immunostaining. (**g**) Quantitative comparison of the median cross-sectional area of muscle fibres in patient and control samples. In (**a**, **b**, **d**, **e**) the median level in each control was set to 1. Dashed lines indicate the normalised median and interquartile range. Data were analysed using a Mann-Whitney test. **** $p$ <0·0001.

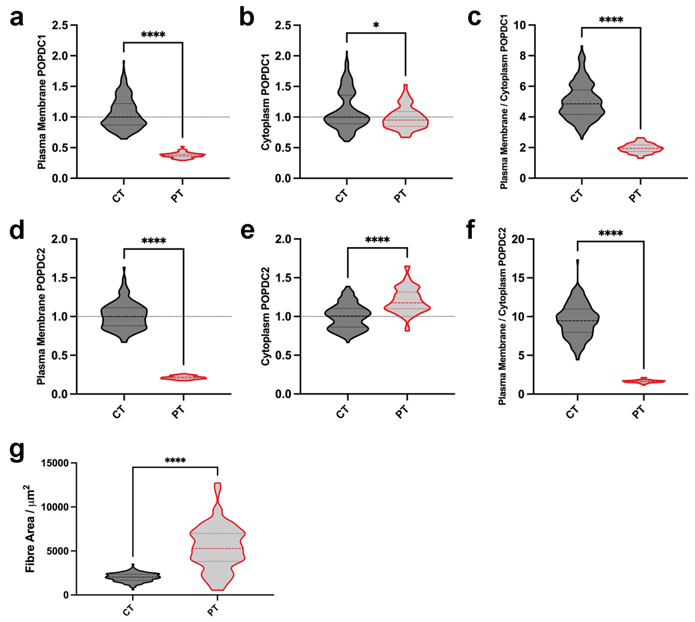


**Supplemental Figure 2. Expression analysis of POPDC1 and POPDC2 in skeletal muscle biopsy of a patient carrying a *POPDC1* p.Q153X mutation.** Expression of (**a**-**c**) POPDC1 and (**d**-**f**) POPDC2 in (**a**, **d**) the sarcolemma, (**b**, **e**) cytoplasm and (**c**, **f**) the ratio of sarcolemma/cytoplasm expression as determined by immunostaining. (**g**) Quantitative comparison of the median cross-sectional area of muscle fibres in patient and control samples. In (**a**, **b**, **d**, **e**) the median level in the control was set to one. Dashed lines indicate the normalised median and interquartile range. Data were analysed using Mann-Whitney test. * $p$ <0·05, **** p <0·0001.

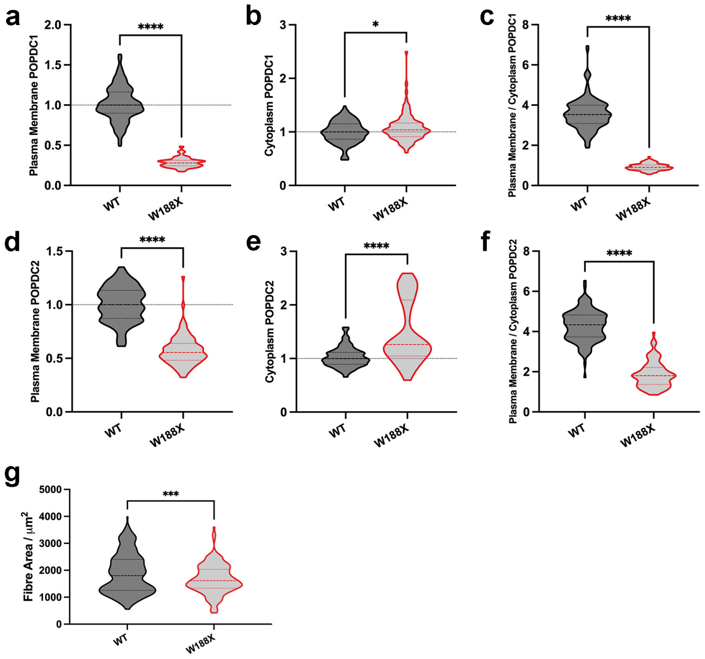


**Supplemental Figure 3. Expression analysis of POPDC1 and POPDC2 in skeletal muscle of homozygous mice carrying a *Popdc2* p.W188X mutation and a WT control.** Expression analysis of (**a**-**c**) POPDC1 and (**d**-**f**) POPDC2 in (**a**, **d**) the sarcolemma, (**b**, **e**) cytoplasm and (**c**, **f**) the ratio of sarcolemma/cytoplasm expression as determined by immunostaining. (**g**) Quantitative comparison of the median cross-sectional area of muscle fibres in mutant and control samples. In (**a**, **b**, **d**, **e**) the median level in the control was set to one. Dashed lines indicate the normalised median and interquartile range. Data were analysed using a Mann-Whitney test. * $p$<0·05, *** $p$ <0·001, **** $p$ <0·0001.

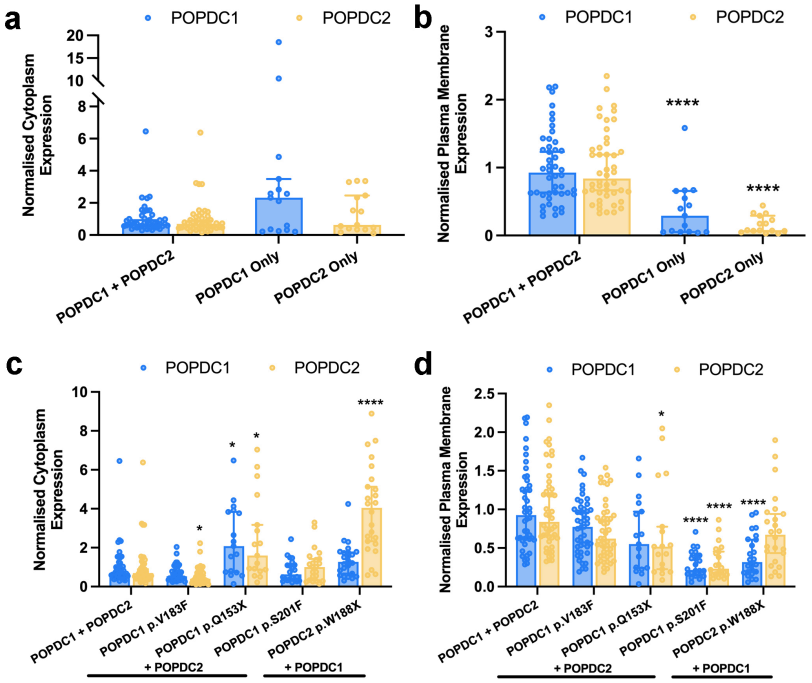


**Supplemental Figure 4. Expression of POPDC1 and POPDC2 in HEK293 cells after transfection with different POPDC1 and POPDC2 constructs. a**) Normalised absolute expression levels of POPDC1-ECFP and POPDC2-EYFP in the cytoplasm and **b**) plasma membrane when co-expressed or expressed alone in HEK293 cells. Bars show median ± 95% CI. Single (POPDC1-ECFP n=9, POPDC2-ECFP n=9) or co-expression conditions (n=46) were compared using a Mann-Whitney test. **c**) Normalised absolute expression levels of POPDC1-ECFP and POPDC2-EYFP in the cytoplasm and **d**) plasma membrane when co-expressed with wildtype (n=46) or mutant (POPDC1 p.V183F n=47, POPDC1 p.Q153X n=17, POPDC1 p.S201F n=22, POPDC2 p.W188X n=24) POPDC dimerization partner in HEK293 cells. Bars show median ± 95% CI. Groups were compared using Kruskal-Wallis followed by Dunn's test using the wildtype pair for comparison; * $p$ <0·05, **** $p$ <0·0001.

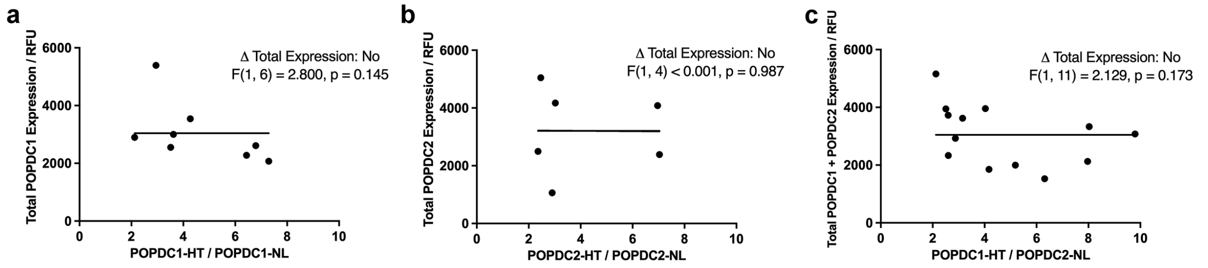


**Supplemental Figure 5. Total BRET expression of POPDC isoforms.** (**a**-**c**) The total expression level of POPDC-NL and POPDC-HT constructs in HEK293 cells when expressed at different ratios of (**a**) POPDC1-NL + POPDC1-HT, (**b**) POPDC2-NL + POPDC2-HT, (**c**) POPDC1-HT + POPDC2-NL. The plots were fitted to a horizontal line and compared to a non-horizontal fit using an F-test with the results shown as insets. Only ratios of POPDC-HT:POPDC-NL expression levels above two were included, as recommended by Felce et al.[1] $p > 0{\cdot}05$ indicates acceptance of a horizontal fit with no significant difference in total POPDC expression at different expression ratios.

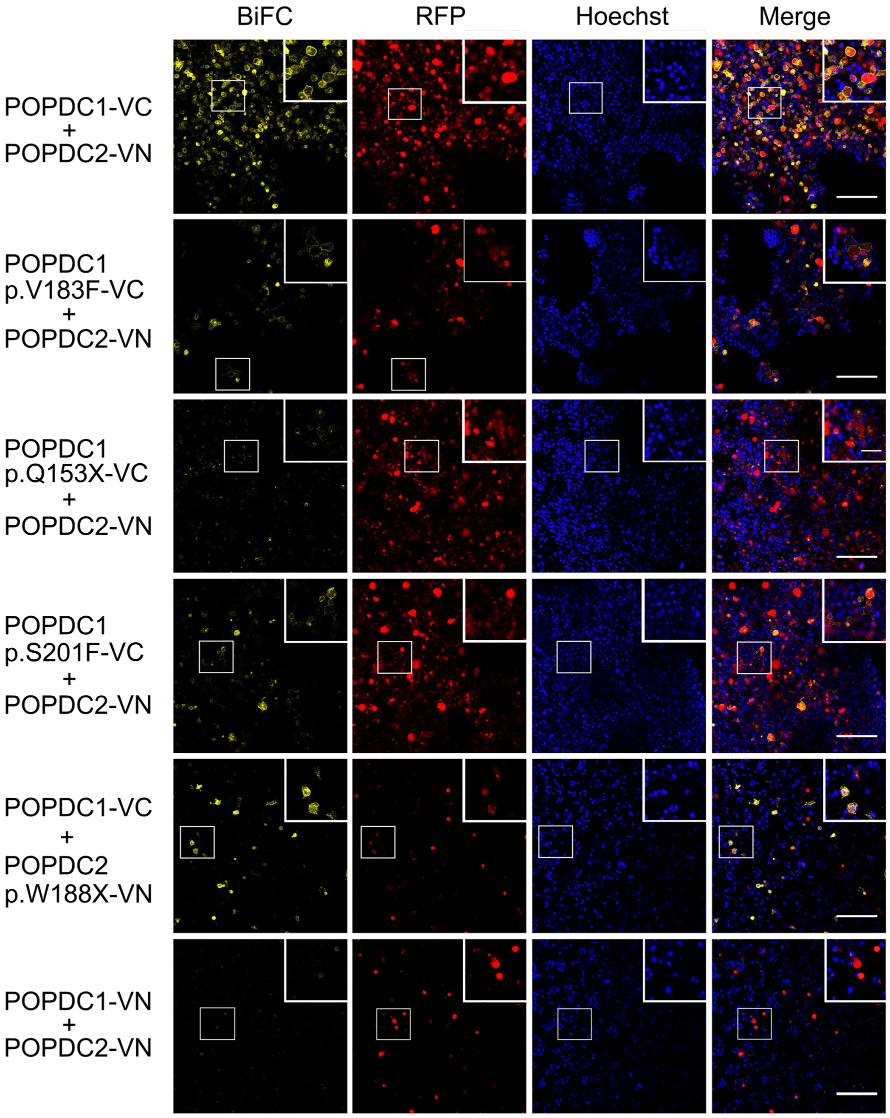


**Supplemental Figure 6. Analysis of bimolecular fluorescence complementation (BiFC) in transfected HEK293 cells.** Confocal microscopy of HEK293 cells after co-transfection of POPDC1-VC or POPDC2-VN with either POPDC1 p.V183F-VC, POPDC1 p.Q153X-VC, POPDC1 p.S201F-VC, POPDC2 p.W188X-VN, respectively. Insets show the boxed area at higher magnification. mRFP was used as internal expression control. The BiFC signal (Venus fluorescence) normalised to mRFP expression was measured only from cells expressing significant levels of mRFP. Nuclei were stained with Hoechst-33342. 5-9 images were taken per group over a minimum of 2 transfections for each construct. Scale bar: 200 μm.

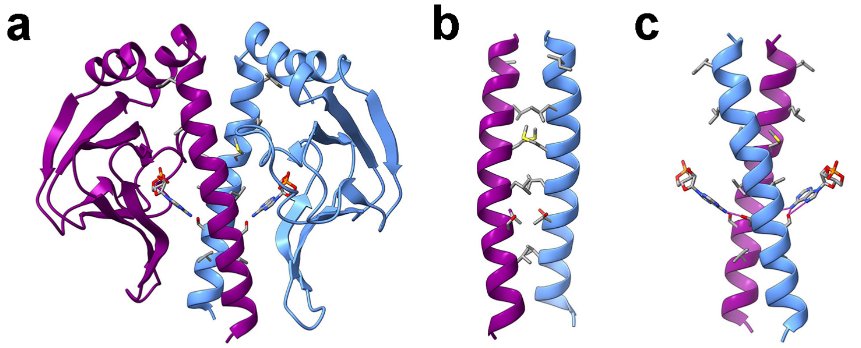


**Supplemental Figure 7. Structural models of CAP protein from E. coli.** (**a**) Structure of a CAP dimer in the cAMP bound form (PDB = 1G6N).[2] (**b**) The αC-helix interface that modulates CAP dimerization. (**c**) Interaction of residues of the αC-helices of CAP with cAMP.

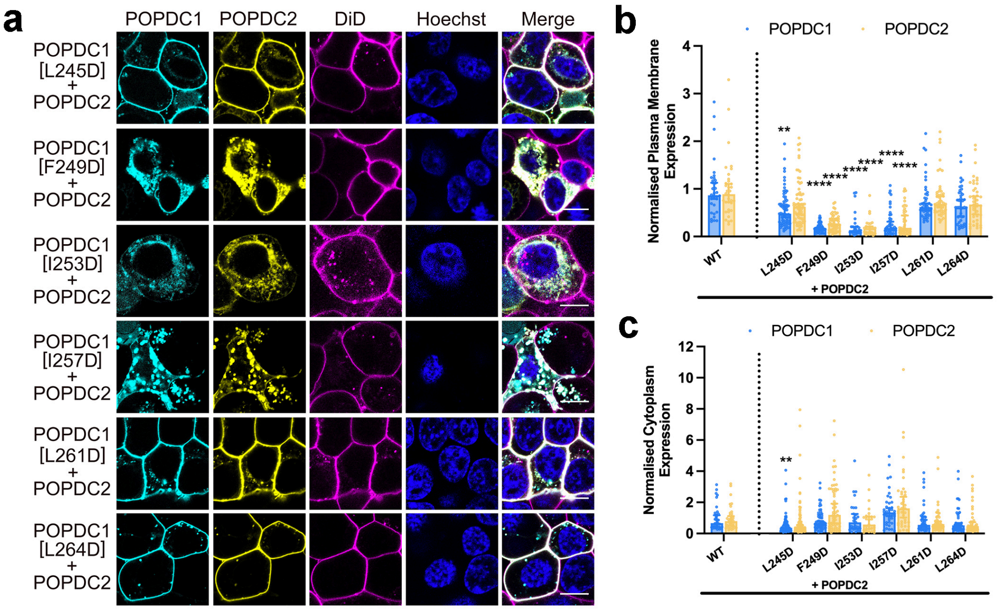


**Supplemental Figure 8. Membrane trafficking after co-transfection of αC-helix mutants of POPDC1 and wildtype POPDC2.** (**a**) POPDC2-EYFP was expressed in HEK293 cells with wildtype POPDC1-ECFP and a series of constructs in which one of the conserved hydrophobic residues within the αC-helix had been mutated to an aspartic acid. Scale bar: 10 µm. (**b**, **c**) Quantification of the normalised expression levels of POPDC1-ECFP and POPDC2-EYFP in the (**b**) plasma membrane and (**c**) cytoplasm as a function of the different POPDC1 mutations. Total number of cells analysed: WT n= 34, L245D n = 71, F249D n = 56, I253D n = 27, I257D n = 40, L261D n = 47, L264D n = 42. Min two transfections per group. Bars show median ± 95% CI. Groups were compared using Kruskal-Wallis followed by Dunn's test using the wildtype pair as a comparison; ** $p$ <0·01, **** $p$ <0·0001.

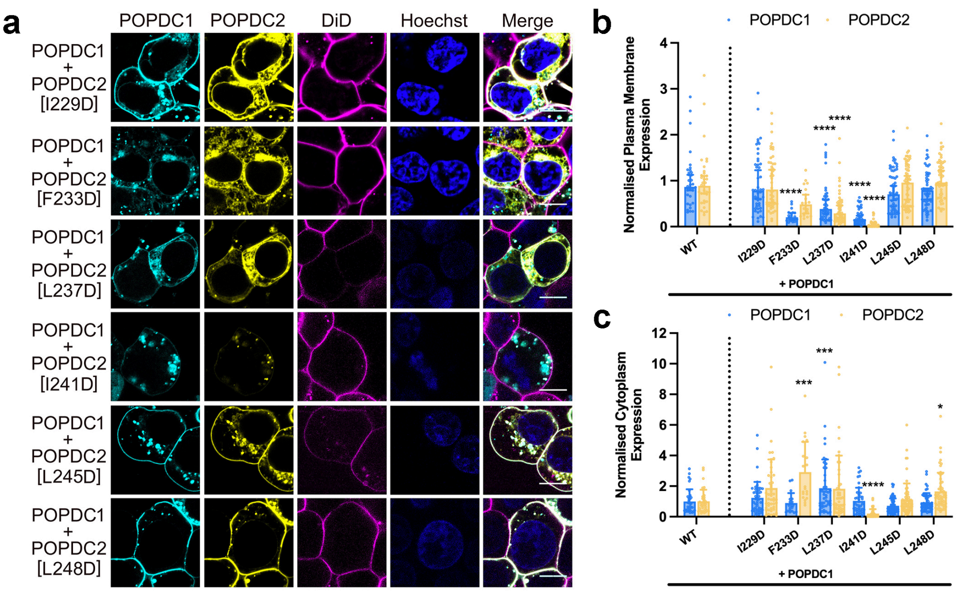


**Supplemental Figure 9. Membrane trafficking after co-transfection of wildtype POPDC1 and αC-helix mutants of POPDC2.** (**a**) POPDC1-ECFP was expressed in HEK293 cells together with wildtype POPDC2-EYFP or a construct in which one of the conserved hydrophobic residues within the αC-helix of POPDC2 had been mutated to an aspartic acid. The plasma membrane was marked using DiD. Scale bar: 10 µm. (**b**, **c**) Quantification of the normalised expression levels of POPDC1-ECFP and POPDC2-EYFP in the (**b**) plasma membrane and (**c**) cytoplasm as a function of the different POPDC2 mutations. Total number of cells analysed: WT n=34, I229D n=48, F233D n=20, L237D n=45, I241D n=42, L245D n=63, L248D n=56. Min 2 transfections per group. Bars show median ± 95% CI. Groups were compared using Kruskal-Wallis followed by Dunn's test using the wildtype pair as a comparison; * $p < 0{\cdot}05$, *** $p < 0{\cdot}005$, **** $p < 0{\cdot}0001$.

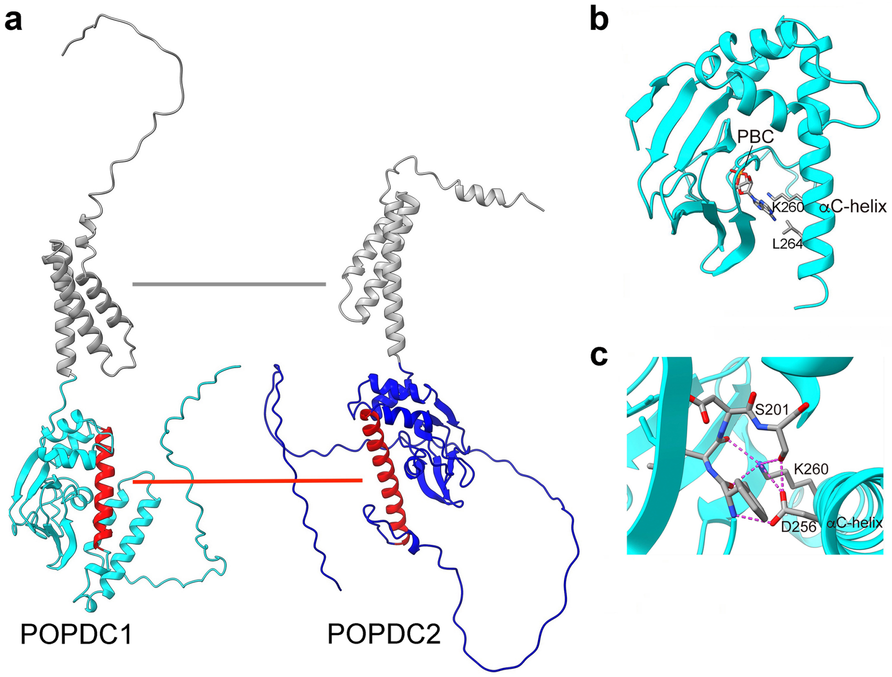


**Supplemental Figure 11. The role of the αC-helix in the function of POPDC proteins.** (**a**) Model of the protein domains mediating the interaction of POPDC1 and POPDC2. The interaction between POPDC1 and POPDC2 is proposed to be mediated through an interface between the αC-helices (red). Another interaction via the N-terminus and/or transmembrane domains (grey) is also depicted. (**b**) The predicted binding mode of cAMP to the Popeye domain of POPDC1, as determined by the 3DLigandSite server. [3] The side chains of K260 and L264 are shown. (**c**) The side chains of the ultra-conserved S201, D256 and K260, in POPDC1 are predicted to form a H-bond network connecting the PBC of the Popeye domain with the αC-helix. Additional H-bonds formed by atoms from the peptide bond are also depicted.

**Supplemental Table 1: Primer sequences used in site-directed mutagenesis of POPDC1-ECFP or POPDC2-EYFP.**

| Template | Mutation | Forward Primer | Reverse Primer |
|---|---|---|---|
| **POPDC1-ECFP** | Q153X | 5'-CGGGATCCACCGGTCGCC-3' | 5'-GATCATGCAAAACTGTCCAGTTAGTCTTCTGAAC-3' |
| | V183F | 5'-AAAAATGAAGTTCTCCTATCGAG-3' | 5'-CCCTTCAAGAGAATACTC-3' |
| | S201F | 5'-TTTATAGATTTTCCTGAATTTAGATCAACTC-3' | 5'-GGCACAGGGGTAAATGTT-3' |
| | L245D | 5’-AGAACCTTTCGACTATGAAATCTTTAGGTATCTTATTG-3’ | 5’-GATTCCAGAAAGTATGTTAATC-3’ |
| | F249D | 5’-GTATGAAATCGACAGGTATCTTATTGGAAAAG-3’ | 5’-AAGAAAGGTTCTGATTCC-3’ |
| | I253D | 5'-TAGGTATCTTGACGGAAAAGACATCACAAATAAG-3' | 5'-AAGATTTCATACAAGAAAGGTTC-3' |
| | I257D | 5'-TGGAAAAGACGACACAAATAAGCTC-3' | 5'-ATAAGATACCTAAAGATTTCATAC-3' |
| | L261D | 5'-CACAAATAAGGACTACTCATTGAATGATC-3' | 5'-ATGTCTTTTCCAATAAGATAC-3' |
| | L264D | 5'-GCTCTACTCAGACAATGATCCCAC-3' | 5'-TTATTTGTGATGTCTTTTCC-3' |
| **POPDC2-EYFP** | W188X | 5'-CGGGATCCACCGGTCGCC-3' | 5'-CTCAGGAGAGTCCATGAACTGGTATGG-3' |
| | I229D | 5’-AGAACCTTTCGACTATGAAATCTTTAGGTATCTTATTG-3’ | 5’-GATTCCAGAAAGTATGTTAATC-3’ |
| | F233D | 5’-GTATGAAATCGACAGGTATCTTATTGGAAAAG-3’ | 5’-AAGAAAGGTTCTGATTCC-3’ |
| | L237D | 5'-TAGGTATCTTGACGGAAAAGACATCACAAATAAG-3' | 5'-AAGATTTCATACAAGAAAGGTTC-3' |
| | I241D | 5'-TGGAAAAGACGACACAAATAAGCTC-3' | 5'-ATAAGATACCTAAAGATTTCATAC-3' |
| | L245D | 5'-CACAAATAAGGACTACTCATTGAATGATC-3' | 5'-ATGTCTTTTCCAATAAGATAC-3' |
| | L248D | 5'-GCTCTACTCAGACAATGATCCCAC-3' | 5'-TTATTTGTGATGTCTTTTCC-3' |
